# Periplasmic protein EipA determines envelope stress resistance and virulence in *Brucella abortus*

**DOI:** 10.1101/442541

**Authors:** Julien Herrou, Jonathan W. Willett, Aretha Fiebig, Lydia M. Varesio, Daniel M. Czyż, Jason X. Cheng, Eveline Ultee, Ariane Briegel, Lance Bigelow, Gyorgy Babnigg, Youngchang Kim, Sean Crosson

## Abstract

Molecular components of the *Brucella abortus* cell envelope play a major role in its ability to infect, colonize and survive inside mammalian host cells. In this study, we have defined a role for a conserved gene of unknown function in *B. abortus* envelope stress resistance and infection. Expression of this gene, which we name *eipA,* is directly activated by the essential cell cycle regulator, CtrA. *eipA* encodes a soluble periplasmic protein that adopts an unusual eight-stranded β-barrel fold. Deletion of *eipA* attenuates replication and survival in macrophage and mouse infection models, and results in sensitivity to treatments that compromise the integrity of the cell envelope. Transposon disruption of genes required for LPS O-polysaccharide biosynthesis is synthetically lethal with *eipA* deletion. This genetic connection between O-polysaccharide and *eipA* is corroborated by our discovery that *eipA* is essential in *Brucella ovis*, a naturally rough species that harbors mutations in several genes required for O-polysaccharide production. Conditional depletion of *eipA* expression in *B. ovis* results in a cell chaining phenotype, providing evidence that *eipA* directly or indirectly influences cell division in *Brucella*. We conclude that EipA is a molecular determinant of *Brucella* virulence that functions to maintain cell envelope integrity and influences cell division.

## Introduction

*Brucella abortus* is a causative agent of brucellosis, a worldwide zoonosis. This bacterium is highly infectious and can be easily transmitted to humans through contact with infected animals and animal products. In humans, disease is often severe and is characterized by multiple sequelae including undulating fever, arthritis, hepatomegaly, splenomegaly, and fatigue. *B. abortus* has the ability to enter and replicate inside mammalian cells (Gorvel & Moreno, 2002), which enables immune evasion and can reduce efficacy of antimicrobial therapies. There are several molecular features of the *B. abortus* cell that play a role in its ability to infect and replicate in mammalian hosts (Atluri *et al.*, 2011, Batut *et al.*, 2004, Byndloss & Tsolis, 2016, Celli, 2006, de Figueiredo *et al.*, 2015, Gorvel, 2008), including smooth lipopolysaccharide (Lapaque *et al.*, 2005, Cardoso *et al.*, 2006, Conde-Alvarez *et al.*, 2012, Smith, 2018), the type IV secretion system (Byndloss & Tsolis, 2016, O’Callaghan *et al.*, 1999, Delrue *et al.*, 2001, den Hartigh *et al.*, 2008), and secreted protein effectors (de Barsy *et al.*, 2011, Myeni *et al.*, 2013, Spera *et al.*, 2013). Here, we report a functional and structural analysis of <u>**e**</u>nvelope <u>**i**</u>ntegrity <u>**p**</u>rotein <u>**A**</u> (EipA), a protein required for *B. abortus* envelope stress resistance and infection.

*Brucella* EipA is a 198-residue protein of unknown function (DUF1134) that has been previously described as one of several dozen conserved “signature proteins” of the class *Alphaproteobacteria* (Figure 1) (Kainth & Gupta, 2005). The promoter region of *eipA* homologs in *Caulobacter crescentus* (gene loci *CC_1035, CCNA_1087*) (Laub *et al.*, 2002, McGrath *et al.*, 2006) and in *Sinorhizobium meliloti* (locus *Smc00651*) (De Nisco *et al.*, 2014, Schluter *et al.*, 2013, Pini *et al.*, 2015) were reported to contain binding motifs for the essential cell cycle regulator CtrA (TTAA-N7-TTAA), which suggested that expression of this protein is tied to the cell cycle. However, functional information on this gene family is limited: *Rhizobium leguminosarum* strains harboring transposon insertions in *eipA* (locus *RL3761*) have a reported growth defect in complex medium (Perry *et al.*, 2016); more recently, a large-scale phenotypic analysis of multiple bacterial species (Price *et al.*, 2018) showed that mutations in *C. crescentus* and *Sphingomonas koreensis eipA* (locus Ga0059261_2034) resulted in antimicrobial susceptibility and a general growth defect in certain defined media, respectively.

**Figure 1:**
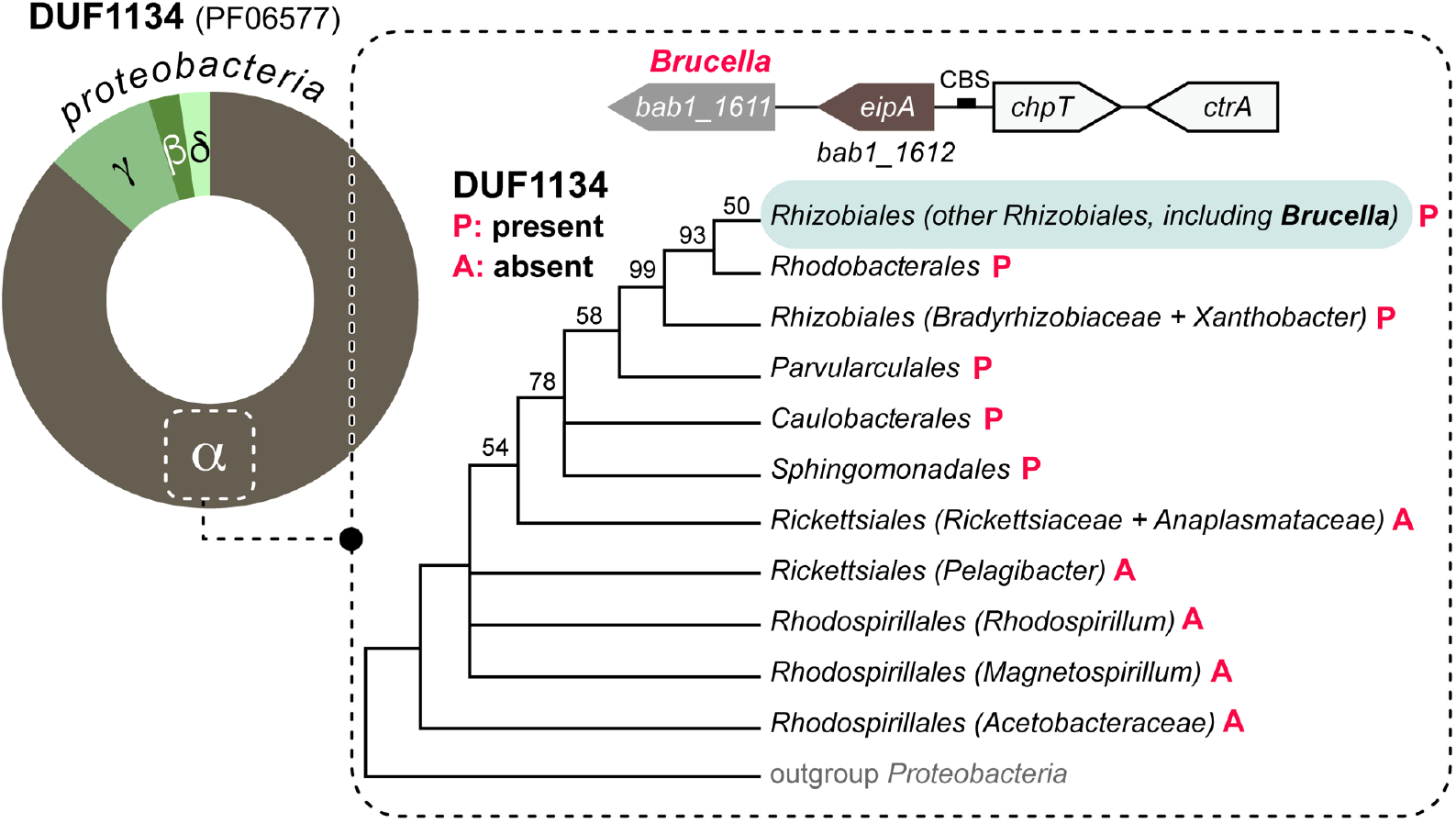
DUF1134 distribution in the bacterial kingdom. Left: DUF1134 is almost entirely restricted to proteobacteria (Finn *et al.*, 2016), and is most common in *Alphaproteobacteria*. The size of the circle slice corresponds to the fractional representation of DUF1134 genes across the *Alpha-, Gamma*-, *Beta*-, and *Deltaproteobacteria*. Right: Bayesian phylogenetic tree showing the distribution of DUF1134 genes in different orders within the class *Alphaproteobacteria* (P= present, A: absent). Bayesian support values are shown when <100%; nodes were collapsed when support was <50%; adapted from Williams *et al*. 2007 (Williams *et al.*, 2007). In *Brucella* spp. (order *Rhizobiales*), *bab1_1612 /* DUF1134 (i.e. *eipA*, in brown) is positioned adjacent to the essential phosphotransferase *chpT* (*bab1_1613*, in white) and the essential response regulator *ctrA* (*bab1_1614*, in white) on chromosome 1. A CtrA-binding site (CBS, black rectangle) is present between *eipA* and *chpT*.

We demonstrate that CtrA does, in fact, directly bind the promoter region of *B. abortus eipA* to activate its expression. EipA folds into a small β-barrel and is secreted to the periplasmic space of the *Brucella* cell. Growth and survival of a *B. abortus* strain in which *eipA* was deleted (D*eipA*) was attenuated in macrophage and mouse infection models. Deletion of *eipA* in *B. abortus* and the related alphaproteobacterium, *C. crescentus*, results in sensitivity to compounds that perturb the integrity of the cell envelope. *eipA* deletion is synthetically lethal with disruption of multiple LPS O-polysaccharide biosynthesis genes in *B. abortus*, and is *eipA* is essential in *Brucella ovis*, a naturally rough species that lacks O-polysaccharide (Tsolis *et al.*, 2009). The results presented herein provide evidence that *eipA* is a molecular determinant of cell envelope integrity in *Alphaproteobacteria*.

## Results

### *eipA* expression is activated by the essential cell cycle regulator, CtrA

*B. abortus* EipA, encoded by gene locus *bab1_1612* (RefSeq: WP_002964697), is a member of sequence family DUF1134 (Bateman *et al.*, 2010) and Pfam PF06577 (Finn *et al.*, 2016). This sequence family is conserved across dozens of genera within the class *Alphaproteobacteria* (Figure 1 and S1). As previously described in *B. abortus* (Willett *et al.*, 2015), *eipA* is co-conserved with the essential cell cycle regulators *chpT* (*bab1_1613*) and *ctrA* (*bab1_1614*), which are also widely-distributed in the *Alphaproteobacteria* (Brilli *et al.*, 2010, Poncin *et al.*, 2018, Panis *et al.*, 2015).

It has been proposed that expression of *eipA* homologs in *C. crescentus* (Laub *et al.*, 2002, McGrath *et al.*, 2006) and in *S. meliloti* (De Nisco *et al.*, 2014, Schluter *et al.*, 2013, Pini *et al.*, 2015) is directly regulated by CtrA, but this prediction has remained untested to date. We sought to determine whether expression of *eipA* is controlled by *B. abortus* CtrA, an established regulator of *Brucella* envelope biology (Francis *et al.*, 2017). The consensus CtrA binding site is TTAA-N7-TTAA (Spencer *et al.*, 2009, Barnett *et al.*, 2001), but the *B. abortus eipA* promoter contains a predicted non-consensus CtrA binding site **TAAA**-(TTCGGGT)-**CTAA**. We conducted an Electrophoretic Mobility Shift Assay (EMSA) with purified CtrA and a ^32^P-labeled DNA oligo corresponding to the promoter sequence of *eipA* (P*eipA*). We observed a full shift of labeled DNA probe in presence of CtrA (Figure 2A). Addition of an excess of equivalent cold (i.e. unlabeled) DNA probe elicited no shift under the same conditions, signifying that the cold probe competes for interaction with CtrA. An excess of a cold DNA probe carrying a mutated CtrA binding site failed to compete CtrA from the specific site (Figure 2A). These results provide evidence that CtrA specifically interacts with the *eipA* promoter region.

**Figure 2:**
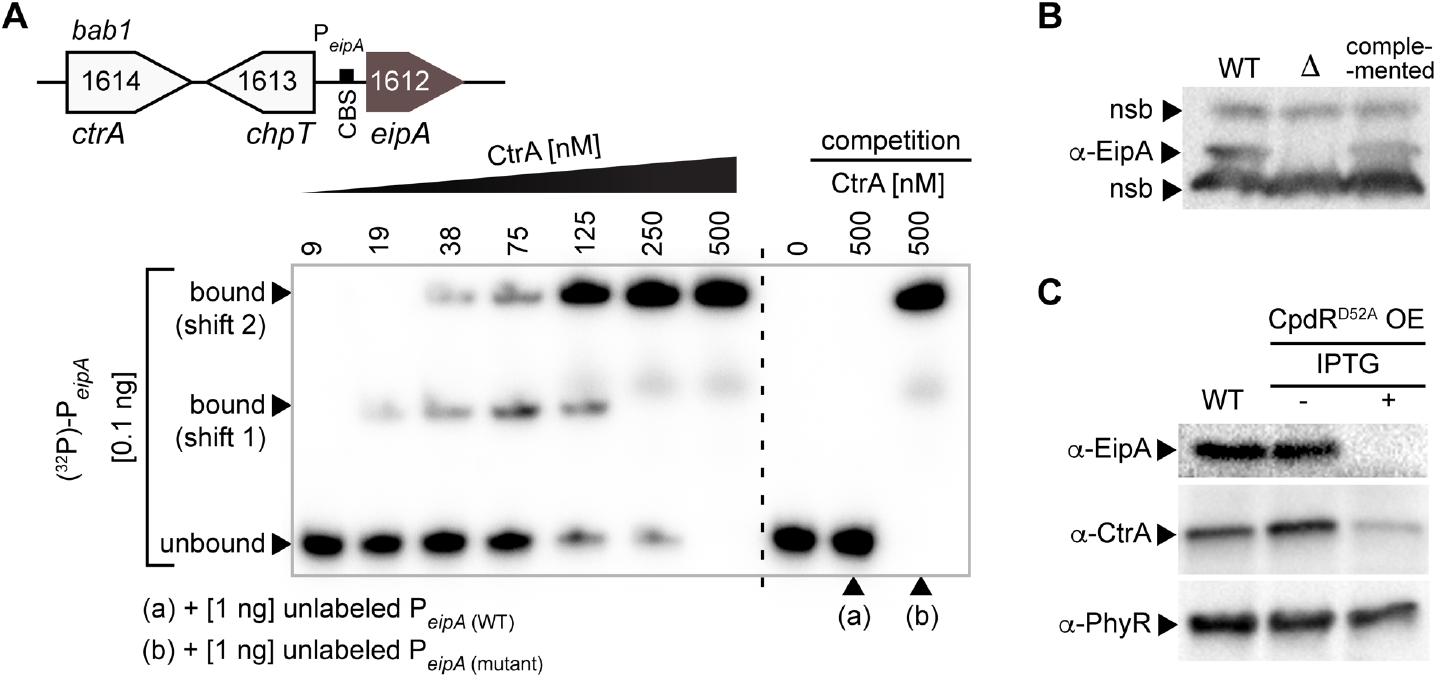
The essential cell cycle regulator, CtrA, directly binds the promoter region of *eipA* in *B. abortus* and activates its expression. A) Electrophoretic mobility shift assay (EMSA) with purified CtrA protein and *eipA* promoter region (P*eipA*). Top: cartoon representation of the *eipA* chromosomal locus, with *ctrA* (*bab1_1614*; white), *chpT* (*bab1_1613*; white), and the CtrA binding site (CBS, black rectangle) present in *eipA* (brown) promoter region. Increasing concentrations of CtrA (9 - 500 nM) were mixed with 0.1 ng of radiolabelled DNA corresponding to *eipA* promoter region (131 bp) (lane 1 to 7). A full shift of the DNA was observed at 500 nM CtrA. Lane 8 shows the DNA alone, without CtrA (0 nM). To test CtrA binding specificity, we competed 0.1 ng of radiolabelled wild-type DNA with 1 ng of unlabelled wild-type DNA (lane 9, (a)) or with 1 ng of unlabeled and mutated DNA (lane 10, (b)). This experiment was independently performed four times; a representative gel is presented. B) Specificity of the rabbit anti-EipA polyclonal serum was tested by western blot using cell lysate from wild-type *B. abortus* (lane 1), the *eipA* deletion strain (lane 2) and the complemented (lane 3) strains. Non-specific bands (nsb) were used as loading controls. C) EipA protein levels were evaluated in wild-type *B. abortus* (lane 1) or in a strain carrying an inducible *cpdR*^D52A^ overexpression (OE) plasmid without (lane 2) or with (lane 3) IPTG inducer. Western blots of lysates from these strains were probed with anti-EipA serum (top), anti-CtrA serum (middle), and anti-PhyR serum as a loading control (bottom). This experiment was independently repeated three times; a representative gel is presented.

Though *eipA* and *chpT* are adjacently positioned on *B. abortus* chromosome 1, and are transcribed from opposite strands (Figure 1), a direct role for CtrA in regulation of *chpT* transcription in *B. abortus* remains untested. The EMSA experiments presented here do not explicitly define a role for CtrA phosphorylation in *eipA* promoter binding in vitro. We note that CtrA phosphorylation is not necessarily required for interaction with the promoter regions of regulated genes, though CtrA phosphorylation is known to increase the affinity of CtrA-DNA interaction (Siam & Marczynski, 2000, Reisenauer *et al.*, 1999, Spencer *et al.*, 2009).

We further tested whether CtrA influenced EipA protein levels in *B. abortus* cells. To measure EipA protein, we developed polyclonal antisera to EipA and confirmed the specificity of this reagent (Figure 2B). As CtrA is essential in *B. abortus* (Willett *et al.*, 2015), we used a conditional mutant to evaluate the effect of CtrA on EipA expression. Specifically, we expressed an inducible variant of CpdR lacking a phosphorylation site (CpdR^D52A^), which activates CtrA proteolysis *in vivo* and thus depletes CtrA protein in the cell (Smith *et al.*, 2014, Willett *et al.*, 2015). As expected, induction of CpdR^D52A^ expression with isopropyl β-D-1-thiogalactopyranoside (IPTG) reduced steady-state CtrA levels in the cell (Figure 2C). This was accompanied by a marked reduction in the steady-state levels of EipA (Figure 2C). These data are consistent with a model in which CtrA binds the *eipA* promoter to activate *eipA* expression.

### *eipA* is required for intracellular replication/survival *in vitro* and for maintenance of spleen colonization *in vivo*

To test whether EipA has role in infection, we infected THP-1 macrophages with wild-type *B. abortus*, ∆*eipA*, and the corresponding complemented Δ*eipA* strains. Infected macrophages were lysed and Colony Forming Units (CFU) were enumerated at 1, 24 and 48 hours post-infection. We observed no significant differences in CFU at the 1-hour time point, indicating that there was no defect in entry for any strain (Figure 3A). At 48 hours post-infection, we observed a significant reduction (≈1.5 log) in the number of intracellular Δ*eipA* cells, providing evidence that *eipA* is required to replicate and/or survive in THP-1 host cells. This defect was fully complemented by restoring the deleted Δ*eipA* locus to wild-type.

**Figure 3:**
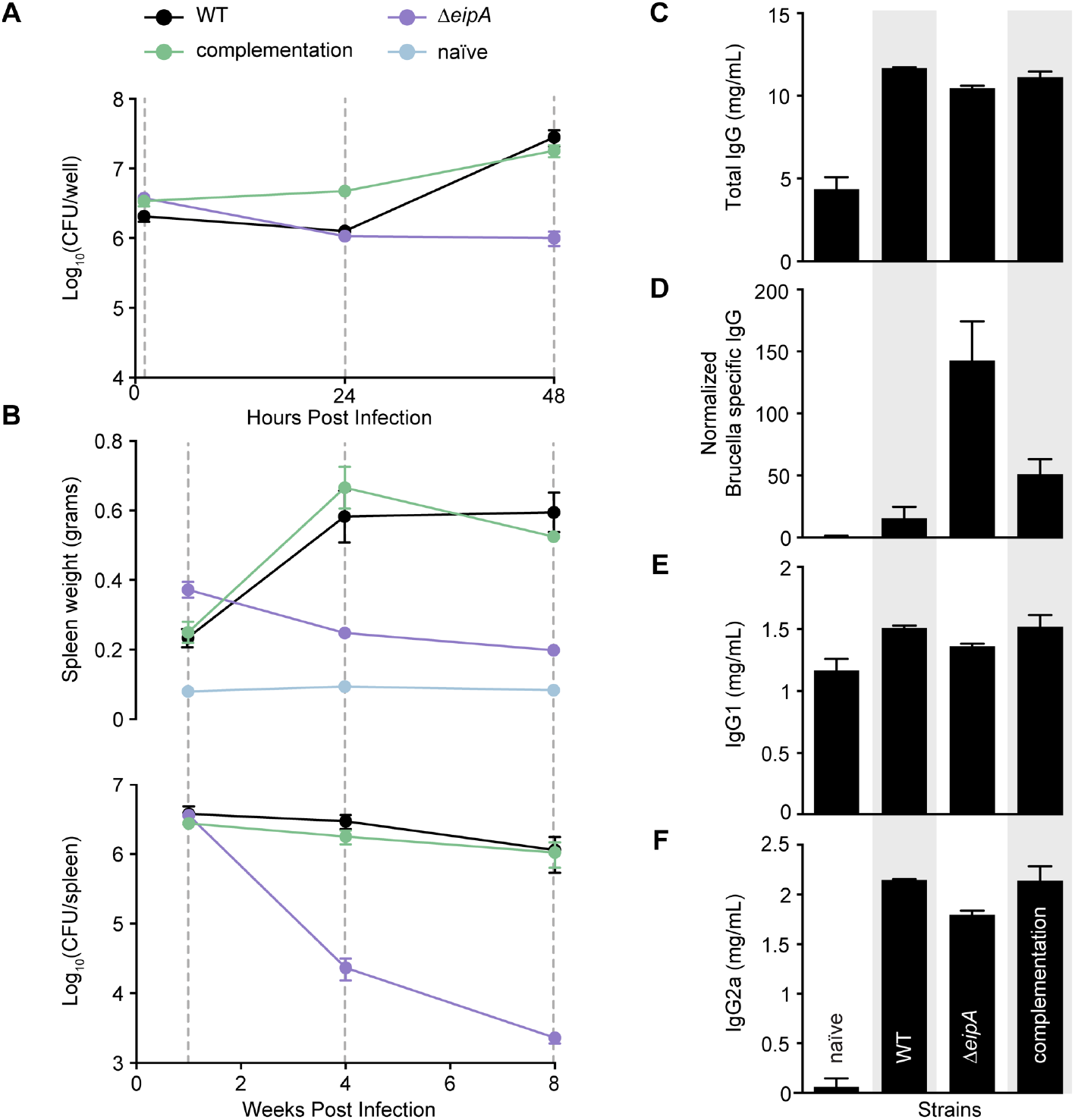
*eipA* is a genetic determinant of *B. abortus* virulence. A) *In vitro* macrophage infection assay: infection of THP-1 cells with wild-type *B. abortus* 2308 (black line), Δ*eipA* (purple line) and the complemented Δ*eipA* strain (green line). The number of *B. abortus* CFUs recovered from THP-1 cells at 1, 24, and 48 hours post-infection is plotted. Each data point (n= 3 per strain) is the mean ± the standard error of the mean. B) *In vivo* mouse infection assay: Female BALB/c mice were injected intraperitoneally with wild-type, Δ*eipA*, or the complemented Δ*eipA* strains. Spleen weights (upper graph) and bacterial burden (lower graph) were measured at 1, 4, and 8 weeks post-infection. Graphs represent data from uninfected naïve mice (light blue) or mice infected with wild-type (black), Δ*eipA* (purple), or the complemented Δ*eipA* (light green) strains. Data presented are the mean ± the standard error of the mean; n= 5 mice per strain per time point. C-F) Antibody quantification in mouse serum harvested at 8 weeks post-infection from naïve control mice or mice infected with wild-type, Δ*eipA*, or the complemented Δ*eipA* strains. Amounts of total IgG (C), *Brucella*-specific IgG (D), IgG1 (E), and IgG2a (F) were determined by ELISA. Each point (naïve: n= 3, WT: n= 2, Δ*eipA* and complemented Δ*eipA*: n= 4) represents the mean ± the standard error of the mean.

We further evaluated the role of *eipA* in a mouse infection model. When compared to BALB/c mice infected with wild-type *B. abortus*, the Δ*eipA* strain exhibited no significant differences in spleen weight or bacterial load at one week post-infection (Figure 3B). At 4 and 8 weeks, mice infected with the wild-type or the complemented deletion strains had pronounced splenomegaly with a load of ≈5 x 10^6^ CFU/spleen. In contrast, mice infected with Δ*eipA* had smaller spleens with reduced numbers of bacteria (Figure 3B). We conclude that *eipA* is not required for spleen colonization but is required for full virulence and chronic persistence in a mouse model of infection.

To assess differences in pathology of mice infected with wild-type and Δ*eipA* strains, we harvested spleens at 8 weeks post-infection. Spleens were fixed, mounted, and subjected to hematoxylin and eosin (H&E) staining (Figure 4). When compared to naïve (uninfected) mice (Figure 4A), spleens of mice infected with wild-type and the complemented Δ*eipA* strains were inflamed, had a significantly reduced white to red pulp ratio due to disruption of lymphoid follicles (follicular lysis) and red pulp expansion, and had increased marginal zones (Figure 4B and D). In addition, we observed higher extramedullary hematopoiesis, histiocytic proliferation, granulomas, and the presence of *Brucella* immunoreactivities (Figure 4B and D). Mice infected with Δ*eipA* had reduced pathologic features with no change in white to red pulp ratio, no follicular lysis or red pulp expansion, and a minimal increase in marginal zones (Figure 4C). Extramedullary hematopoiesis was moderately increased, histiocytic proliferation was minimal, and granulomas and *Brucella* immunoreactivities were rare (Figure 4C). These results are again consistent with a model in which *eipA* is required for full *B. abortus* virulence in a mouse model of infection. A summary of spleen pathology scores is presented in Table S1.

**Figure 4:**
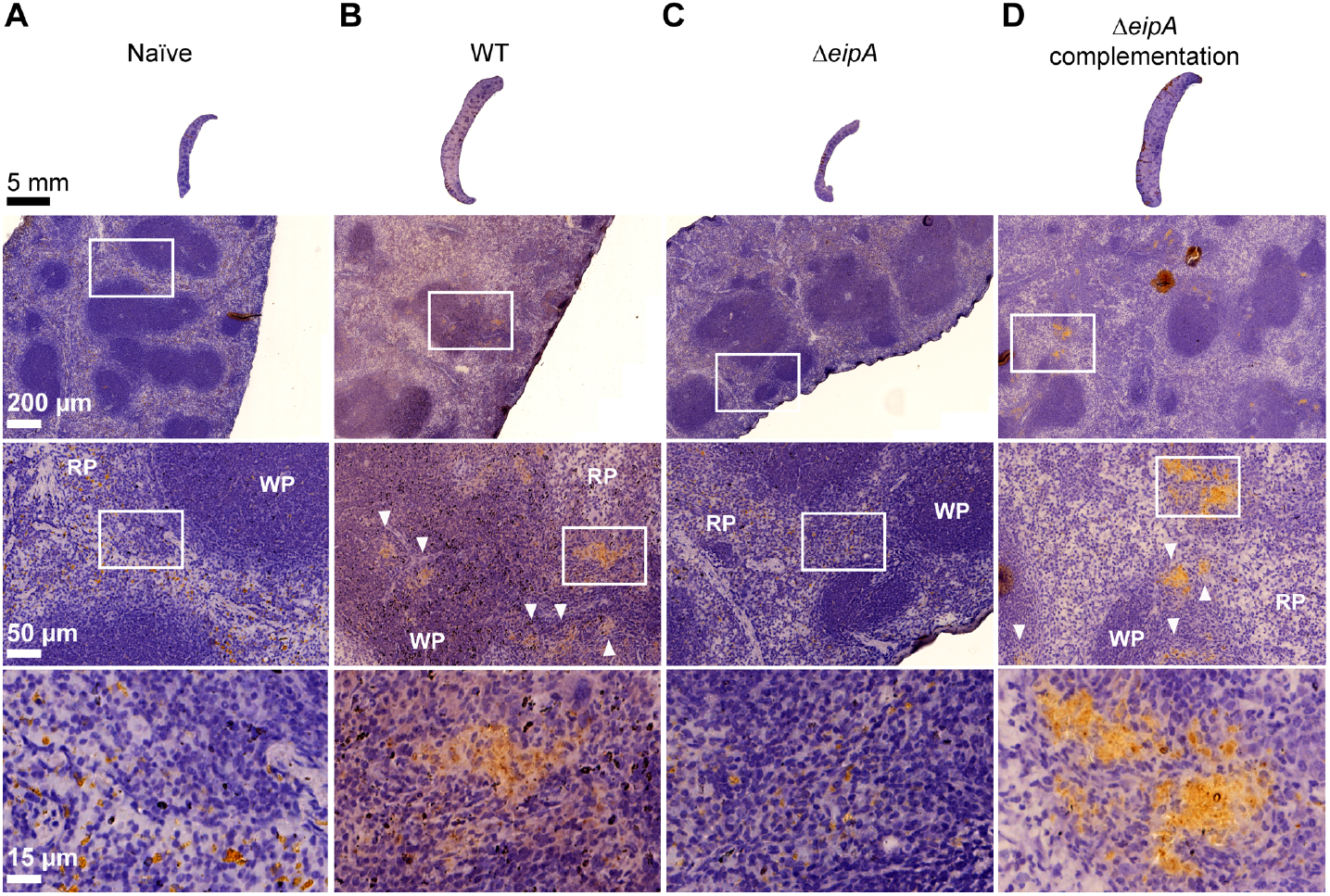
Histology of BALB/c mouse spleens at 8 weeks post-infection. Spleens (n= 1 per strain) were harvested, fixed, and stained with hematoxylin and eosin. Images of spleens from an uninfected mouse (A), and mice infected with wild-type *B. abortus* (B), *B. abortus* Δ*eipA* (C), and the complemented Δ*eipA* strain (D) are presented; white boxes represent specific regions that are shown at increased magnification below each spleen. *Brucella* antigen was visualized by immunohistochemistry with an anti-*Brucella* antibody (brown regions, highlighted with white arrow heads). WP= white pulp, RP= red pulp.

We further measured antibody responses in mice infected with Δ*eipA* and wild-type strains. Serum levels of total IgG, Brucella-specific IgG, IgG1, and IgG2a were measured by enzyme-linked immunosorbent assays (ELISA) (Figure 3C-F). At 8 weeks post-infection, total serum IgG was higher in all infected mice relative to the uninfected control (Figure 3C). The level of *Brucella*specific IgG was ≈3 to 6 times higher in Δ*eipA*-infected mice than in mice infected with the Δ*eipA* complemented or the wild-type strain, respectively (Figure 3D). The wild-type, Δ*eipA* and Δ*eipA* complemented strain-infected mice displayed similar increases in IgG1 and IgG2a at 8 weeks post-infection (Figure 3E and F). We conclude that Δ*eipA* elicits a more robust humoral response than wild-type *B. abortus*. However, we cannot discern whether clearance of the Δ*eipA* strain is mediated by these antibodies, or if antibody production is simply a consequence of antigen release triggered by host clearance of Δ*eipA* by other innate or cell-mediated mechanisms known to control *Brucella* spp. infection in mice (Baldwin & Parent, 2002, Oliveira *et al.*, 2012, Pei *et al.*, 2012, Weiss *et al.*, 2005, Byndloss & Tsolis, 2016).

### *eipA* deletion results in sensitivity to cell envelope stressors

To test whether reduced virulence of the Δ*eipA* strain correlates with sensitivity to particular chemical treatments or growth conditions, we used the Biolog phenotyping platform (Bochner, 2009) to assay growth of wild-type *B. abortus* and the Δ*eipA* strain under 1,920 distinct conditions. The Δ*eipA* strain had growth defects in a number of conditions compared to wild type, including in the presence of molecules known to alter cell envelope integrity. Here forward, we refer to such treatments as “membrane stress” or “envelope stress” (Figure 5A and Table S2). Carbenicillin (plate PM14A – condition G6), an antibiotic that inhibits cell wall synthesis (Bush, 2018), had the strongest effects on Δ*eipA* growth. Captan (plate PM20B – condition G1) and lidocaine (plate PM18C – conditions D11 and D12) also strongly inhibited growth of Δ*eipA*. Captan has been reported to alter membrane integrity in *Saccharomyces cerevisiae* (Scariot *et al.*, 2017) while lidocaine is reported to disrupt bacterial inner membrane potential and permeabilize the outer membrane (Ohsuka *et al.*, 1994). Sodium lactate (plate PM9 – conditions F3 and F4) also impaired Δ*eipA* growth: lactic acid has the capacity to disrupt the Gram negative outer membrane (Alakomi *et al.*, 2000). *B. abortus* Δ*eipA* was also highly sensitive to rolitetracycline (plate PM13B – condition D11), guanine (plate PM13B – condition F6) and dichlofluanid (plate PM16A – condition C3).

**Figure 5:**
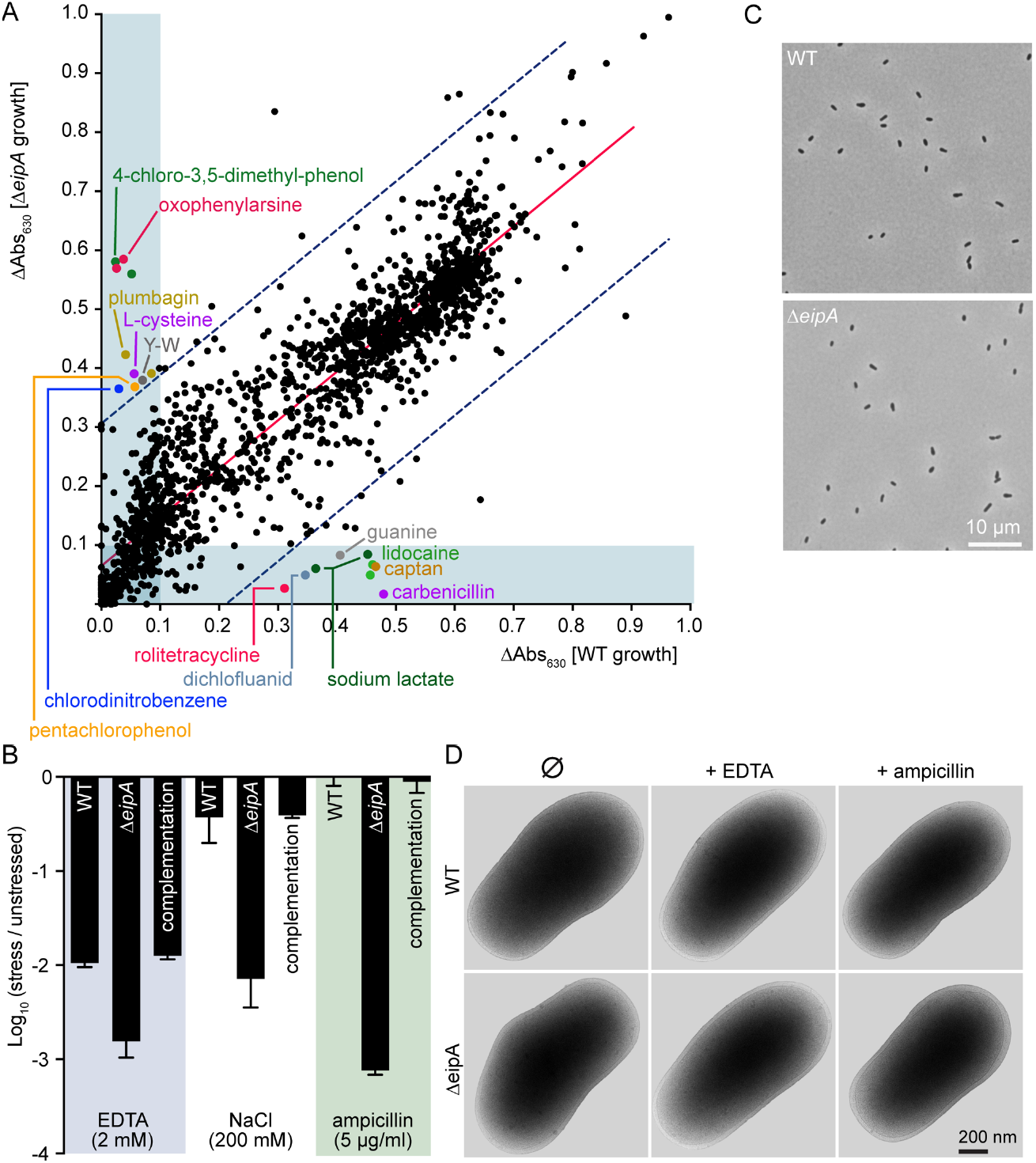
Assessing the effect of cell envelope stressors on *B. abortus* Δ*eipA* growth and survival. A) A high-throughput measurement of Δ*eipA* phenotypes using Biolog phenotype microarrays. Growth of the *ΔeipA* strain (measured as a function of tetrazolium reduction at 630 nm at T=118 hours) plotted against wild-type growth in 1,920 chemical conditions. Red line shows a linear regression fit; the blue dotted lines show the 99% prediction band. Spots outside this band and with Abs630 greater than 0.1 (light blue rectangles) were considered positive hits; demonstrated false positive hits were removed. Concentrations of the different molecules present in the Biolog screen are not available from the manufacturer; this experiment was performed once. B) Validation of select Biolog phenotypes: envelope stress survival assays. Serially diluted cultures of *B. abortus* wild-type, Δ*eipA*, and the complemented Δ*eipA* strains were spotted on plain TSA plates or TSA plates containing EDTA (2 mM), NaCl (200 mM), or ampicillin (5 μg ml^-1^). After 3 days of growth at 37°C / 5% CO2, CFUs for each condition were enumerated and plotted. This experiment was repeated four times; each data point is the mean ± the standard error of the mean. C) Light micrograph of *B. abortus* wild-type (top) and Δ*eipA* (bottom) liquid cultures grown overnight in Brucella broth. D) CryoEM images of *B. abortus* wild-type and Δ*eipA* cells cultivated in liquid broth that either remained untreated or were treated with 2 mM EDTA or 5 μg ml^-1^ ampicillin for 4 hours.

Notably, the Δ*eipA* strain had a growth advantage compared to wild-type in the presence of certain molecules (Figure 5A) including 4-chloro-3,5-dimethyl-phenol (chloroxylenol) (plate PM16A – conditions H7 and H8). This antiseptic is reported to destabilize the cell wall of Gram positive and Gram negative bacteria (McDonnell & Russell, 1999). Δ*eipA* had a relative advantage in pentachlorophenol (plate PM18C – condition C10), which can destabilize membranes in mammalian cells (Duxbury & Thompson, 1987) and in oxophenylarsine (plate PM17A – conditions H11 and H12), a membrane-permeable inhibitor of tyrosine phosphatases (Oetken *et al.*, 1992), which may influence capsular and extracellular polysaccharide synthesis (Standish & Morona, 2014, Heneberg, 2012). *B. abortus* Δ*eipA* also had a small growth advantage in presence of plumbagin (plate PM18C – condition H11 and H12), chlorodinitrobenzene (plate PM16A – condition D3), L-cysteine (plate PM3B – condition A11), and tyrosine-tryptophan dipeptide (plate PM7 – condition H1).

The Biolog assay provided evidence that the Δ*eipA* strain had a cell envelope integrity defect. To further test this hypothesis, we titered wild-type and Δ*eipA* on TSA (Triptic Soy Agar) plates supplemented with known membrane/envelope stressors including EDTA, and ampicillin. We also titered wild-type and Δ*eipA* strains on hyperosmotic (high NaCl) TSA plates. Δ*eipA* had 10 to 1000 times fewer colonies than wild-type on TSA plates containing these compounds (Figure 5B). Restoration of the *ΔeipA* locus to wild-type complemented these defects (Figure 5B). Together, these results provide evidence that *eipA* plays a role in *B. abortus* cell envelope integrity and is required for full resistance to envelope/membrane stressors. Though Δ*eipA* strains are sensitive to these stressors on TSA plates, we observed no apparent morphological defects in liquid broth by light microscopy or cryo-electron microscopy (Figure 5C and D).

As *eipA* is conserved in *Alphaproteobacteria*, we tested whether deleting this gene in another species would result in similar stress sensitivity. It was previously reported that deletion of *C. crescentus eipA* (*CC_1035* / *CCNA_01087*) did not affect cell morphology or growth (McGrath *et al.*, 2006). We tested whether a *C. crescentus* Δ*eipA* strain was sensitive to sodium dodecyl sulfate (SDS), another known membrane stressor. *C. crescentus* Δ*eipA* had a ≈5-log growth/survival defect in the presence of 0.003% SDS compared to wild-type, confirming a role for EipA in cell envelope stress resistance in this species (Figure S2).

### *eipA* is essential in *Brucella ovis*; deletion of *eipA* is synthetically lethal with disruption of genes that function in O-polysaccharide biosynthesis in *B. abortus*

Multiple attempts to delete the *eipA* ortholog in *Brucella ovis* (gene locus *bov_1540;* 100% identity to *bab1_1612*) were unsuccessful, suggesting that *eipA* is essential in this species. We generated and sequenced a *B. ovis* transposon (Tn-Himar) insertion library, which provided further evidence of the essentiality of this gene: no transposon insertions were observed in *eipA* in *B. ovis*. In contrast, we observed many insertions in *eipA* in a *B. abortus* Tn-Himar library that was constructed in parallel (Figure 6A).

**Figure 6:**
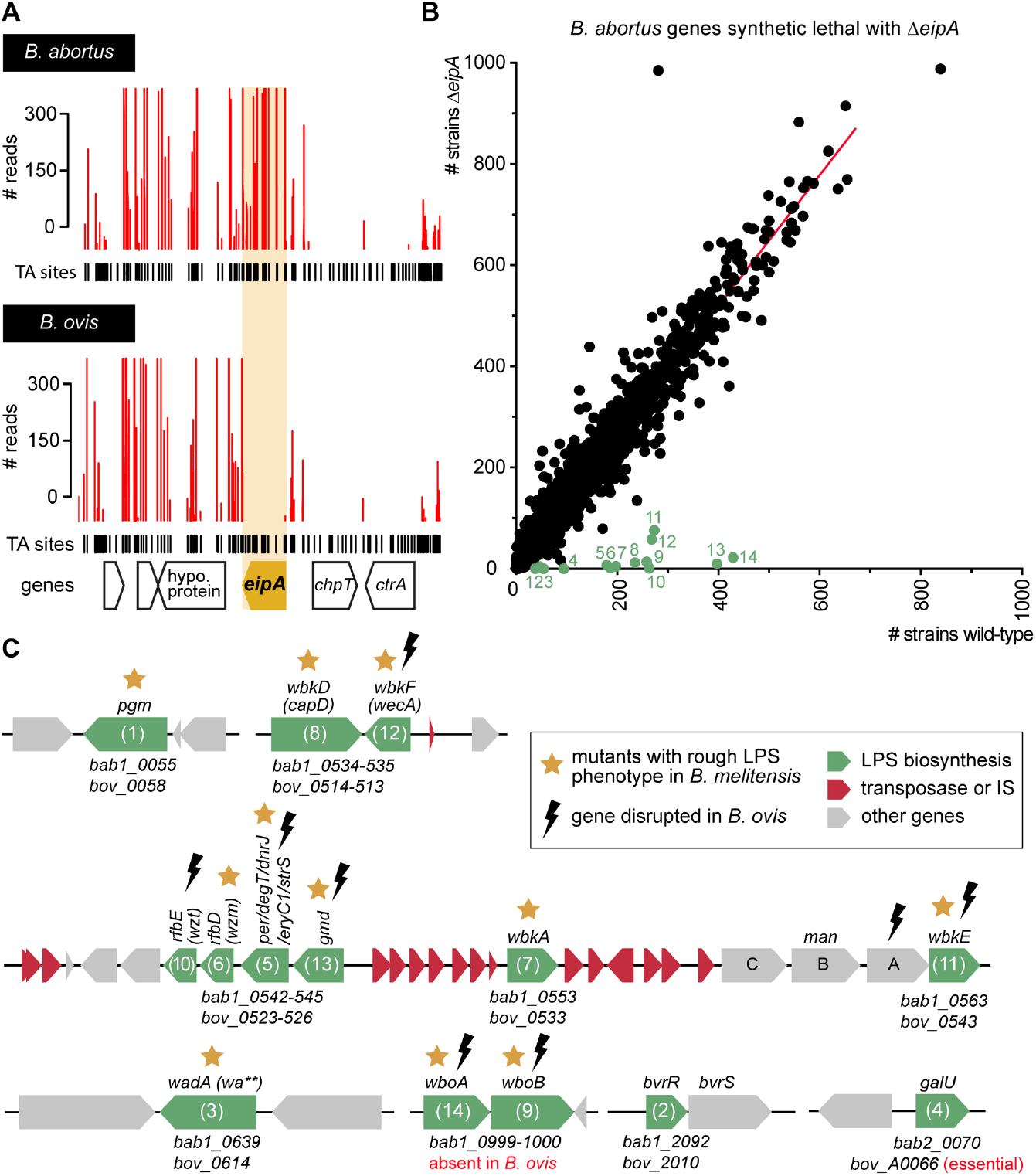
*B. abortus eipA* deletion is synthetically lethal with Tn-disruption of genes involved in LPS O-antigen synthesis; *eipA* is essential in *B. ovis* A) Normalized Tn-Himar insertion profile in the *eipA* region of wild-type *B. abortus* (top) and wild-type *B. ovis* (bottom) chromosome 1. Insertions in the essential genes *chpT* and *ctrA* are not recovered in either species. Insertions in *eipA* are tolerated and recovered in *B. abortus*, but not in *B. ovis*. B) Tn-seq identification of *B. abortus* genes that are synthetically lethal with *eipA.* Unique barcoded Tn insertions (i.e. strains) per gene (black dots) in the *B. abortus* Δ*eipA* Tn-Himar library are plotted as a function of insertions per gene in the wild-type *B. abortus* Tn-Himar library. Synthetic lethal genes (i.e. those genes for which we observed significantly fewer insertions in Δ*eipA* than in wild-type) are represented as green dots and numbered from 1 to 14; the corresponding genes are described in panel C. The red line corresponds to a linear regression. C) Cartoon representation of the synthetic lethal genes presented in panel B. Gene names and locus numbers in *B. abortus* and *B. ovis* are reported above and below the diagram, respectively. Mutations present in these genes that have been reported to confer a rough phenotype in *B. melitensis* are marked with a yellow star. Genes disrupted (by an early stop codon, frame-shift, or mutation) or absent in *B. ovis* are marked by a black lightning bolt. *B. abortus* genes that are synthetically lethal with *eipA* are: *bab1_0055* (*pgm*: phosphoglucomutase), *bab1_0534* (*wbkD/capD*: epimerase/dehydratase), and *bab1_0535* (*wbkF/wecA*: undecaprenyl-glycosyltransferase), *bab1_0542* (*wzt/rfbE*: ATP-binding protein, O-antigen export), *bab1_0543* (*wzm/rfbD*: permease, O-antigen export), *bab1_0544* (*per/degT/dnrJ/eryC1/strS*: perosamine synthetase), *bab1_0545* (*gmd*: GDP-mannose 4,6 dehydratase), *bab1_0553* (*wbkA*: glycosyltransferase), *bab1_0563* (*wbkE*: glycosyltransferase), *bab1_0639* (*wa**/wadA*: glycosyltransferase), *bab1_0999* (*wboA*: glycosyltransferase), *bab1_1000* (*wboB*: glycosyltransferase), *bab1_2092* (*bvrR*: response regulator), and *bab2_0070* (*galU*: UTP-glucose-1-phosphate uridylyltransferase).

To better understand the role of *eipA* in *Brucella* biology, we created a third transposon library in the *B. abortus* Δ*eipA* background aimed at identifying genes that are synthetically lethal with *eipA* deletion. Specifically, we sought to identify genes that are dispensable in wild-type cells, but essential in the absence of *eipA*. We discovered 14 such synthetic lethal genes in *B. abortus* (Figure 6B and C) that have been previously reported to play a role in O-polysaccharide synthesis (Gonzalez *et al.*, 2008, Cardoso *et al.*, 2006, Ugalde *et al.*, 2000, Zhang *et al.*, 2013, Tian *et al.*, 2014, Mancilla *et al.*, 2012, Cloeckaert *et al.*, 2000, Dabral *et al.*, 2015, Monreal *et al.*, 2003, Guzman-Verri *et al.*, 2002, Lamontagne *et al.*, 2007, Manterola *et al.*, 2005, Salvador-Bescos *et al.*, 2018); mutations in this gene set result in a rough phenotype in *Brucella melitensis* (Gonzalez *et al.*, 2008). It is well established that *B. ovis* is a stably rough strain that lacks LPS O-polysaccharide (Mancilla, 2015, Soler-Llorens *et al.*, 2014, Tsolis *et al.*, 2009, Gonzalez *et al.*, 2008). Indeed, many of the O-polysaccharide genes that are synthetically lethal with *eipA* in *B. abortus* are frame-shifted, carry nonsense mutations, or are absent from the genome of *B. ovis* (Figure 6C) (Tsolis *et al.*, 2009, Zygmunt *et al.*, 2009). The degradation of O-polysaccharide genes in *B. ovis* is likely the reason that *eipA* is essential in this species. Notably, *B. abortus* gene *bab1_2092* (*bvrR*) was also synthetically lethal with *eipA* (Figure 6B and C). This response regulator is part of a two-component system (BvrR/S) that contributes to homeostasis of the outer membrane, and regulates LPS and the expression of periplasmic and outer membrane proteins (Guzman-Verri *et al.*, 2002, Lamontagne *et al.*, 2007, Manterola *et al.*, 2005).

### *B. abortus* Δ***eipA* is not a rough strain**

Our data provide evidence for a genetic interaction between *eipA* and O-polysaccharide biosynthesis in *Brucella* spp. Considering that smooth LPS (containing O-polysaccharide) is an important virulence determinant, a loss of smooth LPS in the *B. abortus* Δ*eipA* strain could explain the virulence defect of this strain. To test whether *eipA* is required for smooth LPS synthesis, we assayed wild-type and Δ*eipA* agglutination in the presence of serum from a *B. abortus*-infected mouse. The serological response to smooth *Brucella* species is primarily to LPS (Palmer & Douglas, 1989, Weynants *et al.*, 1996, Cardoso *et al.*, 2006), and thus agglutination provides an indirect indication of whether the LPS O-polysaccharide is present (smooth) or absent (rough). Both *B. abortus* strains agglutinated in the presence of serum, suggesting the presence of O-polysaccharide in Δ*eipA* (Figure 7A). As a negative control, we incubated the stably rough species, *B. ovis*, with the mouse serum; as expected this strain did not agglutinate (Figure 7A). As a second test, we assayed the ability of acriflavine to agglutinate *B. abortus* wild-type and Δ*eipA* strains. Acriflavine is a molecule known to agglutinate rough strains such as *B. ovis*. After two hours of incubation, we observed no agglutination of *B. abortus* wild-type or Δ*eipA* strains. As a positive control, we treated *B. ovis* with acriflavine and observed agglutination (Figure 7B). Together, these data provide evidence that deletion of *eipA* does not result in loss of smooth LPS to an extent that is sufficient to alter agglutination phenotypes.

**Figure 7:**
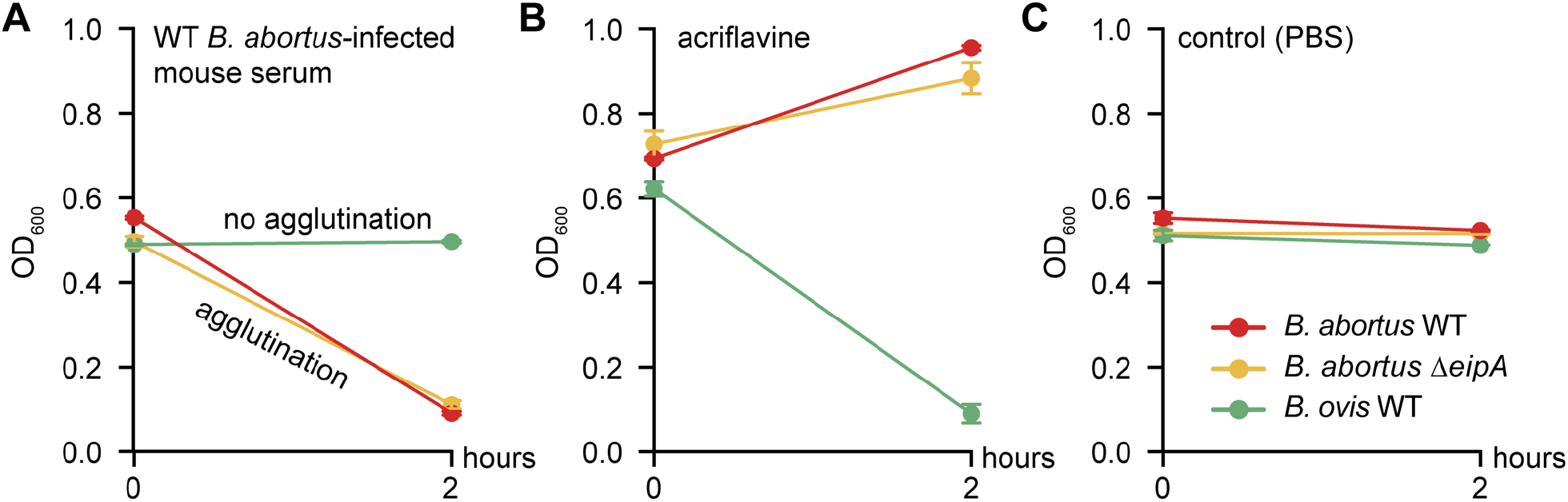
Deletion of *eipA* does not affect the agglutination phenotype of *B. abortus*. Strains (wildtype *B. abortus* in red; Δ*eipA* in yellow) and the wild-type *B. ovis* strain (in green) were incubated for 2 hours at room temperature in 1 ml of PBS supplemented with either (A) 20 μl of serum from a wild-type *B. abortus*-infected mouse, (B) 5 mM acriflavine (final concentration), or (C) no treatment. OD600 was measured for each treatment at the beginning and the end of the experiment (starting OD600 ≈0.5). This experiment was performed in triplicate; each data point is the mean ± the standard error of the mean.

### **EipA is an unusual** β**-barrel protein that is secreted to the periplasm**

Atomic-level 3D structure can provide insight into the molecular basis of protein function. As such, we solved an x-ray crystal structure of *B. abortus* EipA (PDB ID code: 5UC0). EipA formed hexagonal crystals of space group P63 (*α*=*b*=113.075 Å, *c*=64.308 Å, *α*=*ß*=90°, *y*=120°) that diffracted to 1.73 Å resolution; we refined this structure to an Rwork= 0.176 and Rfree= 0.208. Crystallographic data and refinement statistics are summarized in Table S4. Two EipA molecules are present in the crystallographic asymmetric unit; these two molecules interact through hydrogen bonds between the main chains of residues present in β-strand 6 (Figure S3A). Extra electron density, which we assigned to two polyethylene glycol (PEG) molecules, four sulfate ions and two chloride ions, is evident in the EipA crystal structure (Figure S4A).

Each EipA monomer (N39-F198) consists of 10 antiparallel β-strands, with β1-β5 and β8-β10 forming a small β-barrel of ≈17 Å diameter. β6 and β7 form an antiparallel β-hairpin loop that is positioned at the surface of this barrel. Four additional α-helices (α1-α4) are also present, and positioned at the external surface of the β-barrel (Figure 8A). The surface of the protein is positively and negatively charged but presents notable hydrophobic patches (Figure S4B). The barrel lumen is filled with hydrophobic side chains. No cavity or empty space is found within the barrel, and ends are obstructed by α3 and α4, two small α-helices present in the loops connecting β4-β5 and β9-β10, respectively. This structure suggests that EipA, in this conformation, cannot accommodate small molecules or ligands in its lumen (Figure S4C). We searched the Dali database (Holm & Laakso, 2016) with the EipA structure, but did not uncover proteins that are clear structural homologs. Structures of *Rhodococcus equi* virulence-associated proteins - VapG, VapD, and VapB, (Okoko *et al.*, 2015) - had the lowest root mean square deviation (rmsd) (≈3), most significant Z-score (≈7), and highest sequence identity (≈10 and 13%) (Table S5 and Figure S5). However, a functional connection between these proteins, if any, is not known.

**Figure 8:**
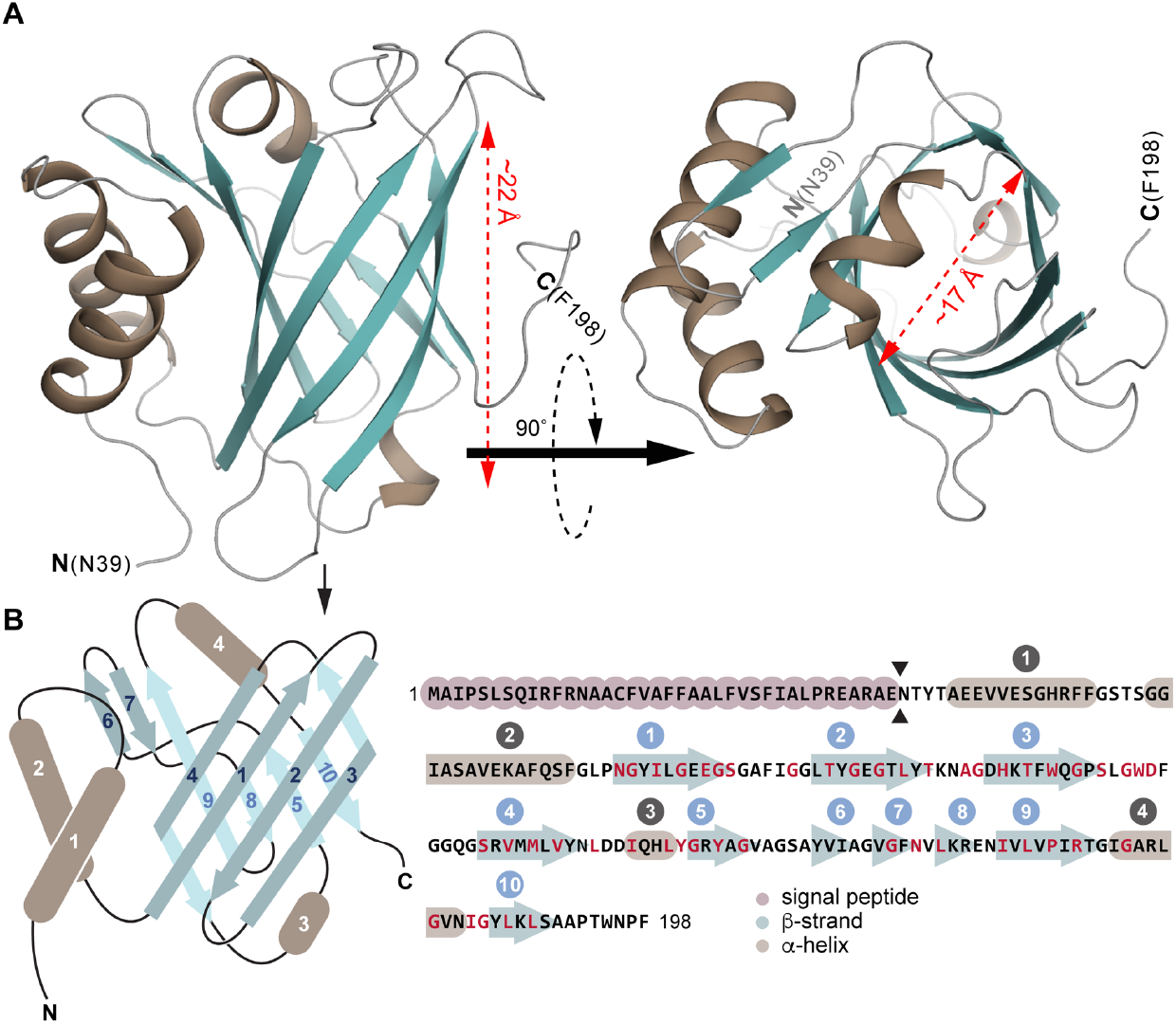
EipA adopts a compact β-barrel structure. A) Left: cartoon representation of the EipA X-ray crystal structure: β-strands are in turquoise and α-helices are in brown. The N-terminus (residue N39) and the C-terminus (residue F198) are annotated. Right: a top view of EipA. Measurements of the β-barrel length and diameter are indicated in red. B) Left: simplified representation of EipA structure with the β-strands in turquoise and α-helices in brown. Eight β- strands (β1, β2, β3, β4, β5, β8, β9, and β10) form a central β-barrel. β6, β7, α1, α2, α3, and α4 adopt an external position at the surface of the β-barrel. Right: amino acid sequence of EipA. The signal peptide sequence (M1-E38) is highlighted in pink. β-strands are in turquoise and α-helices are in brown; secondary structures are numbered above the sequence. Residues with side chains oriented in the lumen of the protein are highlighted in red (74% of these residues are hydrophobic).

EipA is dimeric in the asymmetric unit and, with other dimers present in adjacent asymmetric units, forms an apparent homohexameric complex (Figure S3B). To test whether EipA forms such a complex in solution, we performed size exclusion chromatography on purified *B. abortus* EipA. EipA eluted at a volume with an apparent molecular mass of ≈12 kDa, which is consistent with a monomer (Figure 9A). There was no evidence of larger oligomers by gel filtration. The difference between the calculated molecular mass of EipA (≈20.5 kDa) and the molecular mass measured by gel filtration may be explained by the compact nature of the EipA protein fold.

**Figure 9:**
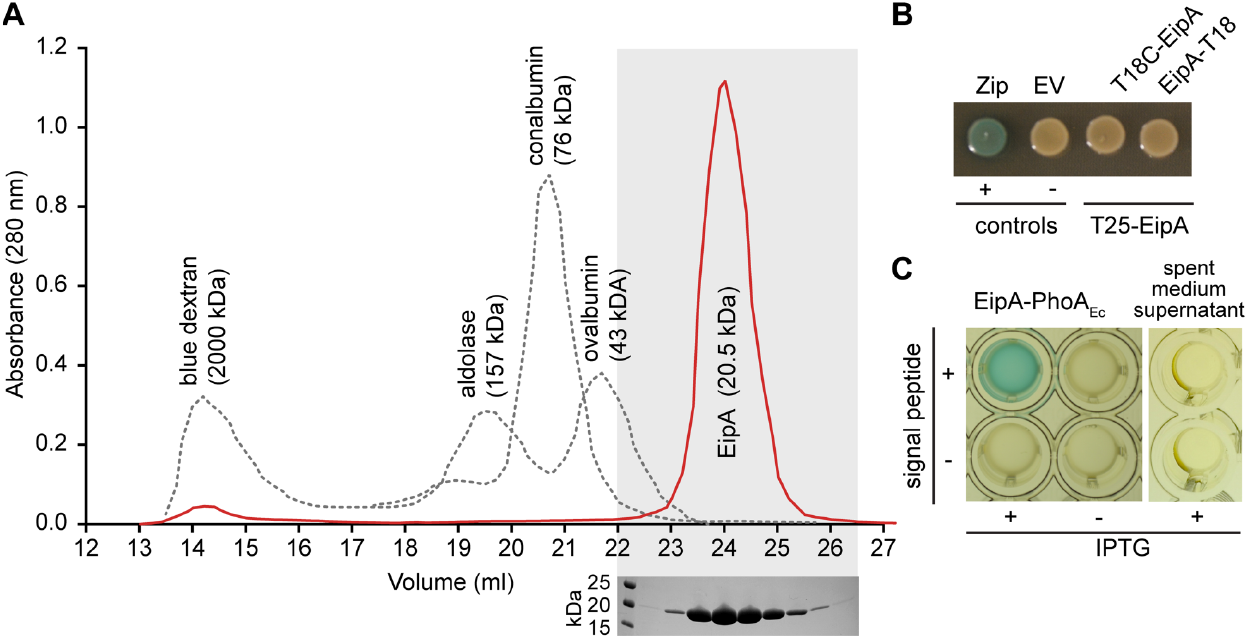
EipA is monomeric in solution and is secreted to the *Brucella* periplasm. A) Top: size exclusion chromatography elution profile of purified EipA (in red). Bottom: the corresponding elution fractions were resolved by SDS-PAGE, and a single ≈20 kDa band was visible on the gel. Elution profiles of the molecular weight standards (blue dextran: 2000 kDa, aldolase: 157 kDa, conalbumin: 76 kDa, ovalbumin: 43 kDa) are shown as a dashed grey line. This experiment was performed twice and yielded similar elution profiles. B) Bacterial two-hybrid assay. EipA fused to the T25 adenylate cyclase fragment was expressed in presence of EipA N-or C-terminally fused to the T18 adenylate cyclase fragment. Leucine-zipper (Zip) and empty vector (EV) strains were used as positive (blue) and negative (beige/yellow) controls, respectively. This experiment was performed at least three times with independent clones. A representative picture of a plate is presented. C) Alkaline phosphatase assay. Overnight cultures of *B. ovis,* expressing EipA with (+) or without (-) its signal peptide and fused to *E. coli* PhoA (PhoAEc), were grown in presence (+) or absence (-) of 1 mM IPTG inducer. In a 96-well plate, these cultures were mixed with BCIP (200 μg ml^-1^ final concentration) and developed for 2 hours at 37°C / 5% CO2. Only the strain expressing EipA-PhoAEc with a signal peptide turned blue, providing evidence that EipA is secreted to the periplasm. As a control, spent medium supernatants were mixed with BCIP to test whether EipA-PhoAEc is secreted from the cell into the medium. After 2 hours of incubation, no color change was observed, indicating that EipA-PhoAEc is not secreted from the cell. These experiments were performed at least three times with independent clones. A representative image is presented.

As a second approach to assess intermolecular interactions between EipA molecules, we performed a bacterial two-hybrid protein interaction assay. EipA^L31-F198^ was fused to the T25, T18 or T18C fragments of the *Bordetella pertussis* adenylate cyclase, and the different combinations (T25-EipA with EipA-T18 or T25-EipA with T18C-EipA) were transformed into *E. coli* BTH101 and spotted on LB agar supplemented with Xgal, kanamycin and carbenicillin. After overnight growth at 30°C, only the positive control showed evidence of protein interaction; the spots corresponding to the different EipA T25/T18 combinations remained white, providing additional evidence that EipA does not form higher order oligomers in cells (Figure 9B).

The N-terminal 38 amino acids (M1-E38) of *Brucella* EipA have signal peptide features, as predicted by SignalP 4.2 (Nielsen, 2017). To evaluate whether EipA is indeed secreted to the periplasm, we created protein fusions to *Escherichia coli* alkaline phosphatase (PhoA) and expressed these fusions from a *lac* promoter in *B. ovis*. Two protein fusions were used: 1) a fusion in which the C-terminus of full-length EipA (M1-F198) was fused to *E. coli* PhoA^R22-K471^ and 2) a fusion that lacked the hypothetical EipA signal peptide (EipA^L31-F198^-PhoA^R22-K471^). After overnight growth in Brucella broth in presence or absence of 1 mM IPTG, we adjusted each culture to the same density and loaded into a 96-well plate containing 5-Bromo-4-Chloro-3-Indolyl Phosphate (BCIP, final concentration: 200 μg ml^-1^). When dephosphorylated by PhoA, BCIP undergoes a color change to blue. Since BCIP diffusion through the inner membrane is inefficient, this reagent can be used to measure PhoA activity in the periplasmic space or in the extracellular medium (Marrichi *et al.*, 2008). After a 2-hour incubation at 37°C, the well containing the *B. ovis* cells expressing the EipA^M1-F198^-PhoA^R22-K471^ fusion turned dark blue. No color change was observed in the well containing the *B. ovis* strain expressing the EipA^L31-F198^-PhoA^R22-K471^ protein fusion (Figure 9C). As expected, lack of induction with 1 mM IPTG resulted in no fusion expression and no color change for either fusion strain (Figure 9C). To test if EipA was secreted into the growth medium, we performed a similar experiment on spent medium supernatants from the different cultures. After 2 hours of incubation, no color change was observed, suggesting that the EipA-PhoA protein fusion is not secreted from the cell into the medium. Our data provide evidence that EipA is a monomeric periplasmic protein that folds into a small, eight-stranded β-barrel with a hydrophobic lumen.

### *eipA* depletion results in a cell division defect in *B. ovis*

*eipA* is essential in *B. ovis* and, therefore, cannot be deleted (Figure 6A). To assess the effects of *eipA* loss in *B. ovis*, we generated a strain in which we placed *eipA* under the control of an IPTG-inducible promoter. In the presence of inducer, we successfully deleted the chromosomal copy of *eipA*. Shifting this strain to Schaedler Blood Agar (SBA) plates lacking IPTG resulted in a dramatic defect in growth/survival relative to wild-type (Figure 10A). This defect was abolished by induction of *eipA* expression with IPTG (Figure 10A). Moreover, the *eipA* depletion strain exhibited a morphological defect in the absence of inducer when cells were cultivated on agar plates: cells did not properly divide resulting in a chaining phenotype (Figure 10B). As expected, addition of IPTG inducer restored morphology to that of the wild-type strain (Figure 10B). This result supports a model in which EipA plays a role in *B. ovis* cell division. It is important to note that we did not observe a defect in growth or morphology by light microscopy when the depletion strain was cultivated in liquid broth without IPTG (Figure S6). The basis of this phenotypic difference between liquid and solid medium is not presently understood.

**Figure 10:**
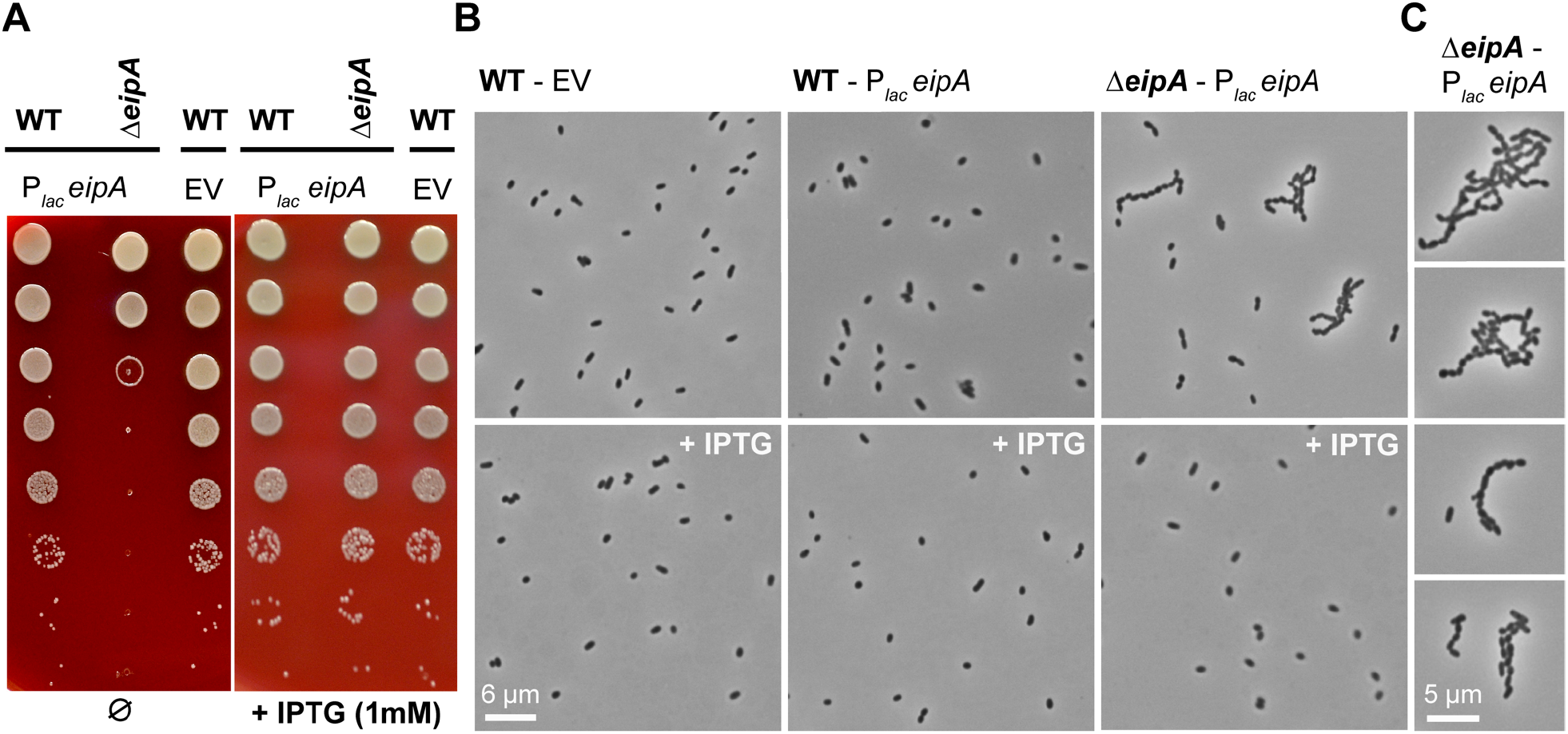
Depletion of *eipA* expression results in a growth defect and a cell chaining phenotype in *B. ovis*. A) Serial dilutions of overnight cultures were spotted on a SBA plates with (right) or without (left) 1 mM IPTG. The *B. ovis* wild-type or Δ*eipA* strains carried a pSRK plasmid (P*lac*-*eipA*) for IPTG-inducible expression of EipA. Wild-type *B. ovis* carrying empty vector (EV) control was also assayed. IPTG-dependent expression of EipA (right) abolishes the growth defect observed in the *B. ovis* Δ*eipA +* P*lac*-*eipA* depletion strain grown without IPTG (left). This experiment was independently performed three times with two different clones each time; a representative picture is presented. B) Light micrographs of the wild-type *B. ovis* empty vector control strain (left), wild-type *B. ovis* + P*lac*-*eipA* strain (middle), and the *B. ovis* Δ*eipA +* P*lac*-*eipA* strain (right) on SBA plates with (bottom) or without (top) 1 mM IPTG. Experiment was performed in triplicate with two independent clones each time; representative pictures are presented for each strain and condition. C) Micrographs highlighting the *B. ovis* cell division/chaining phenotype that is observed upon *eipA* depletion when cells are cultivated on solid media without I

## Discussion

CtrA is an essential regulator of cell development and cell cycle progression in *Alphaproteobacteria* (Brilli *et al.*, 2010). A gene encoding a protein domain of unknown function, DUF1134, is a core member of the CtrA regulon (Laub *et al.*, 2002, McGrath *et al.*, 2006, De Nisco *et al.*, 2014, Francis *et al.*, 2017, Pini *et al.*, 2015, Schluter *et al.*, 2013). The presence of this gene in the CtrA regulon across species strongly suggested that it plays an important role in the biology of *Alphaproteobacteria*. In this study, we have characterized the function of the *B. abortus* DUF1134 protein, which we have named EipA. Expression of EipA is activated by CtrA (Figure 2) and deletion of *eipA* reduced *B. abortus* virulence in both *in vitro* and *in vivo* infection models (Figure 3). A *B. abortus* Δ*eipA* strain is highly sensitive to envelope stressors relative to wild-type, but D*eipA* has no apparent defect in growth or morphology in the absence of stress (Figure 5). *eipA* deletion in *C. crescentus* results in hyper-sensitivity to SDS, providing evidence that the function of this protein is conserved across genera (Figure S2).

Strikingly, *eipA* is essential in the ovine pathogen, *Brucella ovis* (Figure 6A). Unlike *B. abortus*, *B. ovis* is a stably rough species that does not synthesize LPS O-polysaccharide (Tsolis *et al.*, 2009). We defined a genetic connection between *eipA* and smooth LPS, demonstrating that disruption of a set of genes involved in O-polysaccharide production are synthetically lethal with *eipA* deletion in *B. abortus* (Figure 6). The majority of these synthetically lethal genes are degenerated or absent in *B. ovis* (Tsolis *et al.*, 2009), which likely explains the essential nature of *eipA* in this species. LPS O-polysaccharide is known to act as a protective layer against host stressors including cationic peptides, reactive oxygen and complement-mediated lysis and is a key molecule for *Brucella* spp. survival and replication in the host (Cardoso *et al.*, 2006, Martinez de Tejada *et al.*, 1995). However, *B. abortus* Δ*eipA* apparently retains its smooth LPS (Figure 7). Thus, the exact molecular/cellular defect in a *B. abortus* Δ*eipA* that results in stress sensitivity remains undefined at present. One possibility is that EipA influences synthesis, recycling or degradation of bactoprenol. Bactoprenol is a carrier molecule for intermediates in the synthesis of O-chain and peptidoglycan and mutations preventing its recycling or synthesis result in a variety of deleterious or toxic effects such as antibiotic sensitivity, morphological and growth defects, LPS O-chain defects, and reduced virulence (Jorgenson & Young, 2016, Tatar *et al.*, 2007, Yuasa *et al.*, 1969, Bhavsar *et al.*, 2001, Rick *et al.*, 2003, Manat *et al.*, 2014). EipA may also directly interact with enzymes involved in synthesis of peptidoglycan and influence the integrity of the cell wall.

We have further shown that EipA is a monomeric periplasmic protein (Figure 9) that adopts an unusual β-barrel fold, loosely related to *R. equi* virulence-associated proteins (VapG, VapD, VapE) (Figure S5). The structure of EipA alone does not provide clear evidence of its molecular function in the *B. abortus* cell. The EipA β-barrel appears to be too short and hydrophilic to be inserted in the inner or outer membranes. We propose that EipA remains in the periplasm where it may be bound to peptidoglycan or to other proteins as part of a complex. We cannot exclude the possibility that the lumen of EipA binds a ligand. However, ligand binding would require substantial conformational rearrangements of EipA (Figure S4C). Despite substantial progress in characterizing EipA (DUF1134) in this present study, the detailed molecular function of this protein family remains unclear. Considering the data presented here, it is most probable that this protein plays a direct or indirect role in cell division and/or envelope biosynthesis. These two processes are intimately linked to *Brucella* spp. virulence and survival in the host intracellular niche (De Bolle *et al.*, 2015).

In synchronized *C. crescentus* cultures, CtrA-dependent expression of EipA (CC_1035 / CCNA_1087) begins in the middle of the cell cycle S phase and terminates in G2 (Laub *et al.*, 2002, Zhou *et al.*, 2015, Schrader *et al.*, 2016, Fang *et al.*, 2013, McGrath *et al.*, 2007, Hottes *et al.*, 2005, Laub *et al.*, 2000); see CauloBrowser (Lasker *et al.*, 2016) for a compilation of cell cycle expression data (http://www.caulobrowser.org). *C. crescentus* cells become predivisional at this developmental stage and begin to invaginate and divide. Given its expression pattern, it is reasonable to predict that EipA is involved in some aspect of envelope synthesis and/or cell division. Such a model is supported by the chaining phenotype observed in the *B. ovis* depletion strain (Figure 10) and the sensitivity of *B. abortus* and *C. crescentus eipA* deletion strains to envelope stressors (Figures 5 and S2). The molecular connection between O-polysaccharide biosynthesis and the *B. ovis* chaining phenotype upon *eipA* depletion remains undefined, though it is known that alteration of LPS structure can influence morphology and division in *C. crescentus* (Cabeen *et al.*, 2010). It has also been reported that teichoic acids, carbohydrate components of Gram-positive envelopes, play a role in the initiation of cell division (Santa Maria *et al.*, 2014). EipA may serve in part as a bridge that links the state of envelope carbohydrates with the cell division apparatus. Future studies aimed developing a more complete understanding of the role of EipA in control of cell division and cell envelope integrity in *Alphaproteobacteria* are ongoing.

## Materials and Methods

All experiments using live *B. abortus* 2308 were performed in Biosafety Level 3 facilities according to United States Centers for Disease Control (CDC) select agent regulations at the University of Chicago Howard Taylor Ricketts Laboratory.

### Chromosomal deletions in *B. abortus, B. ovis* and in *C. crescentus*

The *B. abortus* Δ*eipA* deletion strain was generated using a double cross-over recombination strategy. Fragments corresponding to ≈500 basepairs 5’ of *eipA*, the first three and last three codons of *eipA*, and ≈500 basepairs 3’ of *eipA* were ligated (T4 DNA ligase, New England Biolabs) into the linearized suicide plasmid pNPTS138, which carries the *nptI* gene for initial selection and the *sacB* gene for counter-selection on sucrose. The ligation was then transformed into *E. coli* Top10 and plated on LB agar plates containing 50 μg ml^-1^ kanamycin. After the sequencing of single colonies, the plasmid was transformed into the wild-type *B. abortus* 2308 strain by conjugation, and single crossover integrants were selected on SBA plates supplemented with kanamycin (50 μg ml^-1^). Counter-selection for the second crossover event was carried out by growing kanamycin-resistant colonies overnight under nonselective conditions and then plating on SBA plates supplemented with 5% sucrose (150 mM). Colonies containing in-frame deletion alleles (encoding a five-amino acid peptide) were identified by PCR screening, and the PCR products were confirmed by sequencing. Genetic complementation of the deletion strain was carried out by transforming the *B. abortus* deletion strain with the pNPTS138 plasmid carrying the wild-type allele locus. For *eipA* deletion in *C. crescentus strain* NA1000, an omega interposon was inserted in the *CC_1035* locus (McGrath *et al.*, 2006). The primers, restriction enzymes, plasmids, and strains used are listed in Tables S6 and S7. To delete *eipA* in a *B. ovis* strain carrying the kanamycin resistant pSRK-*eipA* plasmid, the same strategy was adopted except that a chloramphenicol resistant version of pNPTS138 was used and all the SBA plates and liquid medium used for growth and selection were supplemented with 1 mM IPTG to induce expression of *eipA*.

### Electrophoretic mobility shift assay (EMSA)

Full-length *ctrA* (residues 1-232) from *B. abortus* was PCR amplified. The primers used for this cloning are listed in Table S6. The resulting PCR product was then digested with NdeI / XhoI restriction enzymes (New England Biolabs) and ligated (T4 DNA ligase, New England Biolabs) into similarly digested pET28a plasmid. The ligation was then transformed into *E. coli* Top10. After plating on LB agar plates containing 50 μg ml^-1^ kanamycin, the resulting clones were sequence verified and transformed into the protein overexpression strain *E. coli* Rosetta(DE3) *plysS* (Table S7). For overexpression of His-tagged CtrA, 2 liters of terrific broth (containing 50 μg ml^-1^ kanamycin) was inoculated with a 100 ml of an overnight culture. The culture was grown at 37°C / 220 rpm to OD600 ≈0.8, chilled in an ice bath for 20 minutes and induced with 0.5 mM IPTG. CtrA was allowed to express overnight at 16°C, before cells were pelleted by centrifugation and stored at −20°C. For purification, a cell pellet was resuspended in 50 ml of resuspension buffer (25 mM Tris (pH 7.6), 125 mM NaCl, 10% glycerol) and lysed by two passages through a LV-1 microfluidizer. The lysate was then clarified by centrifugation and mixed with 2.5 ml of Ni-NTA resin on a gravity column. The resin was washed with 5 column volumes (CV) of resuspension buffer, followed by a 5 CV wash with a “detergent” buffer (25 mM Tris (pH 7.6), 125 mM NaCl, 10% glycerol, 5% Triton-X100), a 5 CV wash with a “high salt” buffer (25 mM Tris (pH 7.6), 650 mM NaCl, 10% glycerol) before elution with an imidazole gradient (0 - 500 mM imidazole in resuspension buffer). The purity of the different fractions was assessed on a 12% SDS-PAGE gel. Eluted protein was then dialyzed overnight against a dialysis buffer (25 mM Tris (pH 7.6), 125 mM NaCl, 10% glycerol, 0.5 mM EDTA), before being snap frozen in liquid nitrogen and stored at - 80°C for later experimentation.

For electrophoretic mobility shift assay, a DNA fragment corresponding to the 131 nucleotides present upstream of *eipA* start codon was PCR amplified and used as a probe (see Table S6 for primer information). The DNA fragment was then end labeled using ^32^P-γ-ATP and T4 polynucleotide kinase (New England Biolabs). For gel-shift assays, 0.1 ng of the radiolabeled probe was mixed with increasing concentrations (9 - 500 nM) of purified CtrA in 1 x binding buffer (25 mM Tris (pH 7.6), 1 mM MgCl2, 1 mM MnCl2, 1 mM CaCl2, 50 mM KCl, 2% glycerol, 0.01 mg ml^-1^ BSA, 0.5 μM EDTA). Binding reactions were allowed to proceed for 30 minutes on ice before being loaded on a 8% Tris Borate EDTA (TBE) polyacrylamide gel, equilibrated with 1 x Tris Acetate EDTA (TAE) running buffer and run at 100 volts for 90 minutes at 4°C. To mutate the putative CtrA binding site present in this 131-nucleotide fragment, a gBlock gene fragment (Integrated DNA Technologies, Coralville, IA) carrying mutations in the putative CtrA binding site was used as a PCR template (see Table S6). To test CtrA binding specificity, we competed 0.1 ng of radiolabeled wild-type DNA with 1 ng of unlabeled wild-type DNA or with 1 ng of unlabeled and mutated DNA.

### Western blotting

For Western blot analysis, cultures of *B. abortus* were grown in Brucella broth overnight in the presence or absence of kanamycin (50 μg ml^-1^) and IPTG (5 mM) as required. For each condition, 5 ml of cultures (at OD600 ≈1) were pelleted and resuspended in 500 μl of 4 x SDS loading buffer supplemented with DNase I (New England Biolabs) before boiling at 95°C for 5 minutes. 15 μl of each sample were then resolved on a 12% SDS PAGE gel and transferred to a Millipore 0.45 μm polyvinylidene difluoride (PVDF) membrane (Millipore) using a Trans-Blot turbo transfer system (Bio-Rad). After transfer, the membrane was washed in Tris-buffered saline - Tween 20 (TBST) supplemented with 5% milk and blocked overnight in fresh milk solution. The membrane was then incubated with 10 ml of TBST + 5% milk with an appropriate dilution of anti-EipA (1:5000), anti-CtrA (1:10000) or anti-PhyR (1:5000) rabbit antibodies for 1 hour at room temperature. After a 30-minute wash with TBST, the membrane was incubated for 1 hour in 10 ml TBST + 5% milk with a 1:5000 dilution of anti-rabbit antibody conjugated to Alexa 488 (Life Technologies) secondary antibodies. After a final 30-minute wash in TBST, we used the Bio-Rad ChemiDoc MP imaging system to visualize the bands and quantify the total amount of protein loaded in the gel.

Primary polyclonal EipA rabbit antiserum was generated by Josman LLC. Primary CtrA polyclonal antiserum was a gift from Dr. Peter Chien (University of Massachusetts, Amherst, MA). *Brucella* anti-PhyR polyclonal antiserum (Kim *et al.*, 2013) was used as a loading control.

### Cell culture and macrophage infection assay

Human monocytic THP-1 cells (ATCC TIB-202) were cultured in Roswell Park Memorial Institute (RPMI) 1640 medium supplemented with 10% (v/v) heat-inactivated fetal bovine serum and 2 mM L-glutamine. Differentiation of the cells into macrophage-like cells was induced by the addition of 40 ng ml^-1^ of phorbol 12-myristate-13-acetate (PMA, Sigma) for 48 hours at 37°C in a 5% CO2 atmosphere. Prior to infection, bacteria were harvested from SBA plates, resuspended in sterile RPMI 1640 medium, and cell densities were adjusted. For infection assays, 5 x 10^4^ THP-1 cells were infected with 5 x 10^6^ *B. abortus* cells to achieve a multiplicity of infection of 1:100 in 96-well plates. To synchronize the infections after the addition of the *Brucella* cells, the plates were centrifuged at 200 x *g* for 5 min. After 30 min of incubation at 37°C in a 5% CO2 atmosphere, the medium was removed and replaced with 200 μl of RPMI 1640 medium supplemented with gentamycin (50 μg ml^-1^) and incubated for 1 hour at 37°C in a 5% CO2 atmosphere to kill extracellular bacteria. To determine the numbers of intracellular bacteria at 1 hour postinfection, the cells were washed once with 200 μl of PBS and lysed with 200 μl of PBS supplemented with 0.1% Triton X-100. The lysate was serially diluted and plated on Tryptic Soy Agar (TSA) plates to enumerate CFU. For longer time points (24 and 48 hours), the RPMI + 50 μg ml^-1^ gentamycin was removed and replaced with RPMI + 25 μg ml^-1^ gentamycin to kill extracellular bacteria released over the time course. At 24 or 48-hours post-infection, cells were washed, lysed and *B. abortus* CFU were enumerated as above. For each strain, three independent replicates per time point were performed.

### Mouse infection assay

All mouse studies were approved by the University of Chicago Institutional Animal Care and Use Committee (IACUC). *B. abortus* strains were harvested from SBA plates, resuspended in sterile PBS and diluted to a final concentration of 5 x 10^5^ CFU ml^-1^. For this experiment, 5 mice per strain and per time point were used. One hundred microliters of each bacterial suspension containing 5 x 10^4^ CFU were injected intraperitoneally into 6-week-old female BALB/c mice (Harlan Laboratories, Inc.). At 1, 4, and 8 weeks post-infection, 5 mice per strain were sacrificed, and spleens were removed for weighing and CFU counting. For bacterial enumeration, each spleen was homogenized in 5 ml of sterile PBS supplemented with 0.1% Triton X-100. Each homogenized spleen solution was then serially diluted, and 10 μl of each dilution were spotted on TSA plates and grown for three days at 37°C in a 5% CO2 atmosphere before being counted. At week 8, blood was also collected by cardiac-puncture and serum from each mouse (3 from naïve mice, 2 from WT-infected mice, 4 from mice infected with Δ*eipA* and the complemented Δ*eipA* strain) was separated from blood cells using a serum separation tube (Sarstedt). Sera were subsequently used for Enzyme-Linked ImmunoSorbent Assays (ELISA).

### Determination of antibody responses at 8 weeks post-infection

Mouse serum Immunoglobulin G (IgG) titers were measured using an ELISA in 96-well ELISA plates (Nunc-immuno MaxiSorb, Sigma). Total mouse serum IgG, IgG1, and IgG2a titers were measured using mouse-specific ELISA kits by following manufacturer’s instructions (eBioscience, ThermoFisher). To determine *Brucella*-specific IgG titers, ELISA plates were coated with 100 μl of heat-killed bacterial suspensions prepared in PBS (OD600=2.0) from *B. abortus* grown for 2 days on SBA plates. Cultures were heat-killed at 65°C for 1 hour, cooled to room temperature, and treated with kanamycin (50 μg ml^-1^) and gentamycin (50 μg ml^-1^) to prevent bacterial growth. The signal from all ELISA plates was measured at 450 nm with a 570 nm background correction using a Tecan Infinite M200 PRO fluorescence plate reader.

### Spleen histology

At 8 weeks post-infection, spleens (n= 1 per strain) were prepared for histology. Spleens were first fixed with formalin for 7 days. Formalin was removed and replaced with fresh formalin and fixed for another 7 days before washing and subsequently transferring spleens to 70% ethanol. Whole spleens were submitted for tissue embedding, Hematoxylin and Eosin (H & E) staining, and immunohistochemistry to Nationwide Histology (Veradale, Washington). For immunohistochemistry, goat anti-*Brucella* IgG was used (Tetracore, Inc). We subsequently analyzed and scored slides at the University of Chicago. Pictures of fixed mouse spleens were taken at the University of Chicago Integrated Light Microscopy Facility on a Cambridge Research and Instrumentation whole slide scanner fitted with an Allied Vision Technologies Stingray F146C color camera.

### Bacterial Phenotype MicroArray screening

Bacterial Phenotype MicroArray (PM) assays were carried out according to manufacturer’s protocols (Biolog). Briefly, *B. abortus* strains (wild-type and Δ*eipA*; strain information available in Table S7) were streaked out and cultivated at 37°C and 5% CO2 for 48 hours on an SBA plate, restreaked, and grown for another 48 hours. Cells were scraped off the plate and resuspended in 1x IF-0a medium (Biolog) at a final density equivalent to 5% transmittance at OD600. PM inoculating fluids (PM1-20) were prepared from 0.2 μm filter-sterilized stock solutions at the following final concentrations: ***PM1-2*:** 2 mM MgCl2, 1 mM CaCl2, 25 μM L-arginine, 50 μM L-glutamic acid, 5 μM β-NAD, 25 μM hypoxanthine, 25 μM 5’-UMP, 25 μM L-cystine (pH 8.5), 0.005% yeast extract, and 0.005% tween 40; ***PM3*, *6*, *7, 8*:** 20 mM tricarballylic acid (pH 7.1), 2 mM MgCl2, 1 mM CaCl2, 5 μM β-NAD, 25 μM hypoxanthine, 25 μM 5’-UMP, 25 μM L-cystine (pH 8.5), 0.005% yeast extract, 0.005% tween 40, 0.625 mM D-glucose, and 5 mM pyruvate; ***PM4*:** 20 mM tricarballylic acid (pH 7.1), 2 mM MgCl2, 1 mM CaCl2, 25 μM L-arginine, 50 μM L-glutamic acid, 0.005% yeast extract, 0.005% tween 40, 0.625 mM D-glucose, and 5 mM pyruvate; ***PM5*:** 20 mM tricarballylic acid (pH 7.1), 2 mM MgCl2, 1 mM CaCl2, 0.625 mM D-glucose, and 5 mM pyruvate; ***PM9-20*:** 2 mM MgCl2, 1 mM CaCl2, 0.005% yeast extract, 0.005% tween 40, 2.5 mM D-glucose, and 5 mM pyruvate. PM1-8 inoculating fluids (IF) were prepared in a final 1x concentration of IF-0a GN/GP (Biolog) while PM9-20 inoculating fluids were prepared at a final 1x concentration of IF-10b GN/GP (Biolog). All solutions had a Dye Mix G containing tetrazolium (Biolog) added to a final 1x concentration. A 1:13.6 dilution of 5% transmittance *B. abortus* suspension was added to each inoculating fluid. 100 μl per well of *B. abortus*-containing inoculating fluids were dispensed to each PM plate. PM plates were incubated at 37°C and 5% CO2, and absorbance of tetrazolium reduced by bacterial growth (at 630 nm) was measured daily for 5 days using a Tecan Infinite 200 PRO microplate reader. We note that the concentrations of the different molecules present in this screen are not available from the manufacturer. Molecules that absorb in the visible region of the spectrum often yield false-positive hits, which we removed.

### Plate stress assays

For stress assays, the *B. abortus* strains were first grown for three days on SBA plates. Cells were then harvested and resuspended in sterile PBS to an OD600 of ≈0.015 (≈1 x 10^8^ CFU ml^-1^). The different cell suspensions were then serially diluted and 3 μl of each dilution were spotted on plain TSA plates or TSA plates supplemented with 5 μg ml^-1^ ampicillin, 200 mM NaCl, or 2 mM EDTA. Plates were grown for three days at 37°C in a 5% CO2 atmosphere before being counted. All experiments were performed in quadruplicate and compared to strains grown on plain TSA plates (no stress). The same protocol was used to test the SDS tolerance *C. crescentus* wild-type and *ΔeipA* strains (see Table S7 for strain information); PYE agar plates with or without 0.003% SDS were used for this assay and grown at 30°C for 3 days.

### Cryo-electron microscopy

Overnight *Brucella* cultures were used to inoculate (at a starting OD600 of ≈0.015) 2 ml of Brucella broth supplemented with or without 2 mM EDTA or ampicillin (5 μg ml^-1^). After 4 hours of treatment at 37°C / 5% CO2 / 220 rpm, cells were harvested by centrifugation (5000 x *g* / 10 min) and the pellets were resuspended in PBS + 4% formaldehyde to fix the cells. Cell killing was confirmed before sample removal for imaging. After 1 hour, cells were pelleted and resuspended in 500 μl EM buffer (20 mM Tris (pH 7.6), 50 mM glucose, 10 mM EDTA). Fixed *Brucella* cells were vitrified on glow-discharged 200 mesh copper EM-grids with extra thick R2/2 holey carbon film (Quantifoil). Per grid, 3 μl of the sample were applied and automatically blotted and plunged into liquid ethane with the Leica EM GP plunge-freezer. Images were collected on a Talos L120C TEM (Thermo Fischer) using the Gatan cryo-TEM (626) holder. The images were acquired at a defocus between 8-10 μm, with a pixel size of 0.458 nm.

### Building and mapping the wild-type *B. abortus*, the *B. abortus* Δ*eipA* and the wild-type *B. ovis* Tn-Himar mutant libraries

To build and map the different Tn-Himar insertion libraries, we used a barcoded transposon mutagenesis strategy (Wetmore *et al.*, 2015). Specifically, the *E. coli* APA752 donor strain (strain WM3064 harboring a pKMW3 mariner transposon vector library) was conjugated with wild-type *B. abortus, B. ovis* or with the *B. abortus* Δ*eipA* strain. The WM3064 strain is a *pir*+, diaminopimelic acid (DAP) auxotroph, competent to transduce plasmids. 100 ml of LB broth + 300 μM DAP were inoculated with a vial of *E. coli* APA752 donor strain and incubated overnight a 37°C / 220 rpm. The next day, the culture was pelleted and mixed with *Brucella* (*ovis* or *abortus*) freshly harvested from 4 SBA plates. The donor and recipient were mixed at an approximate ratio of 1:10 based on optical density. The mixture was then spotted on a SBA plate supplemented with 300 μM DAP and incubated a 37°C / 5% CO2 overnight. The next day, the mating was resuspended in 25 ml of Brucella broth and OD600 was measured (OD600 ≈16). The Himar transposon inserts into TA dinucleotide sites in the genome; *Brucella* has ≈78,000 TA sites in its genome when considering the sequence of both strands (10 - 15% are considered to be essential), therefore, approximately 350,000 to 750,000 unique clones will yield a library with 5 - 10 x coverage at each site. To achieve this library diversity, the cell suspension was diluted to OD600 ≈0.016 in 25 ml of Brucella broth, and 1 ml was plated on 25 large (150 mm diameter) SBA plates supplemented with kanamycin (50 μg ml^-1^) to yield approximately 20,000 - 40,000 colonies per plate. After three days of growth at 37°C / 5% CO2, the colonies from all 25 plates were harvested and resuspended in 25 ml of Brucella broth, the OD was measured (OD600 ≈95) and this cell suspension was used to inoculate 500 ml of Brucella broth with kanamycin (50 μg ml^-1^) at OD_600_=0.15. After 6 hours of growth at 37°C, the culture was pelleted and resuspended to an OD600 of ≈0.4 in 25 ml of Brucella broth containing 25% glycerol. 1 ml aliquots were frozen and stored at −80°C until needed.

An aliquot of each Tn insertion library was used to extract the genomic DNA for insertion site mapping. Cells were pelleted (1 min at 12,000 rpm) and resuspended in 100 μl of a Tris EDTA (TE) buffer (10 mM Tris (pH 8), 100 μM EDTA). 500 μl of GES lysis buffer were added (for 100 ml of GES lysis solution: 60 g Guanidinium thiocyanate, 20 ml of a 500 mM EDTA (pH 8) solution, 5 ml of 10% v/v lauryl sarkosyl, completed with water). The solution was vortexed and incubated at 60°C for 15 minutes; 250 μl of cold 7.5 M ammonium sulfate were then added. The solution was vortexed and incubated on ice for 10 minutes. 500 μl of 24:1 chloroform:isoamyl alcohol were added, vortexed, and centrifuged at 12,000 rpm for 15 minutes. The upper aqueous phase (≈700 μl) was transferred to a new tube and 600 - 700 μl of cold 100% isopropanol were added to precipitate the genomic DNA. After mixing, the DNA was pelleted at 12,000 rpm for one minute and washed with 70% ethanol. After removing any residual ethanol, the genomic DNA was resuspended in 50 μl of TE buffer.

To build an Illumina sequencing library enriched in sequences containing the transposon junction, we followed the method of Wetmore *et al.* (Wetmore *et al.*, 2015). Custom adapters were generated by annealing complementary oligos (Mod2_TS_Univ: 5’-ACGCTCTTCCGATC*T-3’, and Mod2_TruSeq: 5’-Phos-GATCGGAAGAGCACACGTCTGAACTCCAGTCA-3’). Both oligonucleotides were diluted in TE buffer to 100 μM and 10 μl of each oligo were mixed. For annealing, the PCR cycling conditions were: 30 minutes at 95°C, and 15 minutes at 70°C. The resulting adapter was diluted to 25 μM using TE buffer. Genomic DNA was sheared to generate 300 - 500 bp fragments. The size selection, end repair, ligation of the custom adapter, and clean up step were performed by the University of Chicago Functional Genomics facility. The PCR amplification step to enrich TN containing fragments was performed using a transposon specific primer, Nspacer_BarSeq_pHIMAR (5’-ATGATACGGCGACCACCGAGATCTACACTCTTTCCCTACACGACGCTCTTCCGATCTNNNNNNCGCCCTGCAAGGGATGTCCACGAG-3’), and an adapter specific primer, P7_MOD_TS_index (5’- CAAGCAGAAGACGGCATACGAGATCGTGATGTGACTGGAGTTCAGACGTGTGCTCTT-3’). Primers with unique indices were used for each sample that was sequenced in the same lane. The following PCR amplification protocol was used: a 100 μl PCR reaction was prepared by mixing 50 μl of GoTaq Green (2x) master mix (Promega) with 10 μl of 50% DMSO, 0.5 μl of each primer (100 μM), 5 μl of sheared adapted-ligated genomic DNA and completed with water and cycled under the following conditions: 94°C for 2 minutes, 25 cycles of [94°C for 30 seconds, 65°C for 20 seconds, 72°C for 30 seconds], 72°C for 10 min. Quality of the PCR product was evaluated on a 2% agarose TBE gel; a smear between 150-500 bp is expected. The PCR products were cleaned with size selection beads to remove fragments less than 200 bp. PCR products were then sequenced with a 150 bp single end Illumina run using standard Illumina sequencing primers. Next, the barcoded transposons were assigned to insertion sites using the perl script MapTnSeq.pl (available at https://bitbucket.org/berkeleylab/feba.git) as previously outlined (Wetmore *et al.*, 2015).

Statistics for the three different transposon insertion libraries are reported in Table S3. For each Himar insertion library, Tn-seq read data have been deposited in the NCBI sequence read archive: *B. abortus* 2308 wild-type (BioProject PRJNA493942; SRR7943723), *B. ovis* ATCC 25840 wild-type (BioProject PRJNA493944; SRR7943724), *B. abortus* Δ*eipA* (Δ*bab1_1612*) (BioProject PRJNA493947; SRR7943771).

### Agglutination assay

To compare the agglutination phenotype of our different strains (wild-type *B. abortus* strain, Δ*eipA* strain, and wild-type *B. ovis*; see Table S7 for strain information), the bacteria were grown on SBA plates for three days at 37°C / 5% CO2, harvested and resuspended in sterile PBS at OD600 ≈0.5. At time (0), 1 ml of each cell suspension was loaded in a spectrophotometer cuvette and mixed with 20 μl of wild-type *B. abortus*-infected mouse serum or with acriflavine (final concentration 5 mM) and ODs were measured at 600 nm. As a control, 1 ml of each cell suspension was also kept in a spectrophotometer cuvette without serum or acriflavine. The different samples were then incubated at room temperature for 2 hours and clarification of the samples, which corresponds to the cells agglutinating, was then measured at OD600. Cuvettes containing PBS supplemented with serum or acriflavine were used as blanks. For each agglutination condition, three independent cell suspensions per strain were tested.

### Protein expression and purification for crystallization

The DNA fragment corresponding to the EipA protein (residues 39 - 198) was PCR amplified. The PCR product was then cloned into pMCSG73 plasmid using a ligation-independent procedure (Aslanidis & de Jong, 1990, Eschenfeldt *et al.*, 2009). The primers used for cloning are listed in Table S6. After transformation of the ligation into *E. coli* Top10 strain, the resulting colonies able to grow on LB agar + ampicillin (100 μg ml^-1^) plates were sequence verified. pMCSG73 plasmids with the correct insert were purified and transformed in the *E. coli* BL21-Gold(DE3) strain for protein expression (see strain information listed in Table S7). The pMCSG73 is a bacterial expression vector harboring a tobacco vein mottling virus-cleavable N-terminal NusA tag and a TEV-cleavable N-terminal His6 and StrepII tag. For protein expression, a 2-liter culture of enriched M9 medium (Kim *et al.*, 2011) (+ 100 μg ml^-1^ ampicillin) was grown at 37°C with shaking at 190 rpm. At OD600 ≈1, the culture was cooled to 4°C and supplemented with 90 mg of L-seleno-L-methionine (Se-Met, Sigma) and 25 mg of each methionine biosynthetic inhibitory amino acid (L-valine, L-isoleucine, L-leucine, L-lysine, L-threonine, and L-phenylalanine). Protein expression was induced overnight at 18°C using 0.5 mM IPTG. After centrifugation, cell pellets were resuspended in 35 ml of lysis buffer (500 mM NaCl, 5% (v/v) glycerol, 50 mM HEPES (pH 8.0), 20 mM imidazole, and 10 mM β-mercaptoethanol) per liter of culture and treated with lysozyme (1 mg ml^-1^). The cell suspension was sonicated, and debris was removed by centrifugation. The Se-Met protein was purified via Ni^2+^-affinity chromatography using the ÄKTAxpress system (GE Healthcare). The column was washed with the lysis buffer and eluted in the same buffer containing 250 mM imidazole. Immediately after purification, the His tag was cleaved at 4°C for 24 to 48 hours using a recombinant His-tagged TEV protease, resulting in an untagged protein with an N-terminal Ser-Asn-Ala peptide. A second Ni^2+^-affinity chromatography purification was performed to remove the protease, non-cleaved protein, and affinity tag. The purified protein was then dialyzed against 20 mM HEPES (pH 8), 250 mM NaCl, and 2 mM DTT buffer. Protein concentrations were determined by UV absorption spectroscopy (280 nm) using the NanoDrop 1000 spectrophotometer (Thermo Fisher Scientific). The purified Se-Met EipA protein was concentrated to 43 mg ml^-1^ using a centrifugal filter (10 kDa MWCO, Amicon-Millipore).

### Crystallization

Initial crystallization screening was carried out using the sitting-drop, vapor-diffusion technique in 96-well CrystalQuick plates (Greiner Bio-one). Trays were prepared using a Mosquito robot (TTP LabTech) and commercial crystallization kits (MCSG-1-4, Anatrace). The drops were prepared by mixing equal volumes (0.4 μl) of the purified protein (43 mg ml^-1^) and the crystallization solution, and equilibrated against 135 μl of the same crystallization solution. Crystallization trays were incubated at 16°C. After 3 days, EipA crystallized in the hexagonal space group P63 from the condition #15 of the MCSG-2 crystallization kit that contains 200 mM ammonium sulfate, 100 mM sodium cacodylate (pH 6.5), 30% PEG8000. Prior to flash freezing in liquid nitrogen, crystals were cryo-protected by briefly washing them in the crystallization solution containing 25% glycerol.

### Crystallographic data collection and data processing

Se-Met crystal diffraction was measured at a temperature of 100 K using a 3-second exposure/degree of oscillation and was collected for 120°. Crystals diffracted to a resolution of 1.73 Å and the corresponding diffraction images were collected on the ADSC Q315r detector with an X-ray wavelength near the selenium edge of 12.66 keV (0.97929 Å) for SAD phasing at the 19-ID beamline (SBC-CAT, Advanced Photon Source, Argonne, Illinois). Diffraction data were processed using the HKL3000 suite (Minor *et al.*, 2006). EipA crystals were twinned and the data had to be reprocessed and scaled from the hexagonal P6322 space group to the lower symmetry space group P63 with the following cell dimensions: *a*=*b*=113.08 Å, *c*=64.31 Å, and *α=β=* 90°, *γ=*120° (see Table S1). The structure was determined by SAD phasing using SHELX C/D/E, mlphare, and dm, and initial automatic protein model building with Buccaneer software, all implemented in the HKL3000 software package (Minor *et al.*, 2006). The initial model was manually adjusted using COOT (Emsley & Cowtan, 2004) and iteratively refined using COOT, PHENIX (Adams *et al.*, 2002), and REFMAC (Murshudov *et al.*, 1997); 5% of the total reflections was kept out of the refinement in both REFMAC and PHENIX throughout the refinement. The final structure converged to an Rwork of 17.6% and Rfree of 20.8% and includes two protein chains (A: residues 39-198; and B: 39-198), 4 sulfates, two chloride ions, two polyethylene glycol (5 C-C-O chains) molecules, and 322 ordered water molecules. The EipA protein contained three N-terminal residues (Ser-Asn-Ala) that remain from the cleaved tag. The stereochemistry of the structure was checked using PROCHECK (Laskowski *et al.*, 1993), and the Ramachandran plot and was validated using the PDB validation server. Coordinates of EipA have been deposited in the PDB (PDB code 5UC0). Crystallographic data and refined model statistics are presented in Table S4. Diffraction images have been uploaded to the SBGrid data server (Data DOI: 10.15785/SBGRID/444).

### Protein sequence alignment and structural homology

Amino acid sequences were aligned using the M-COFFEE Multiple Sequence Alignment Server (Wallace *et al.*, 2006) and shaded using BoxShade. Figures of the structures, structural alignments, electrostatic potential representations and r.m.s.d. calculations were performed using PyMOL (PyMOL Molecular Graphics System, version 1.7.4; Schrödinger, LLC). Surface hydrophobicity was evaluated using the YRB python script (Hagemans *et al.*, 2015). The XtalPred server (Slabinski *et al.*, 2007) and Dali server (Holm & Rosenstrom, 2010) were used to identify proteins with the highest structural and sequence homology. The BLAST server (https://blast.ncbi.nlm.nih.gov/Blast.cgi) was used to identify homologs of *B. abortus* EipA in different taxa within the *Alphaproteobacteria*.

### Size exclusion chromatography

A DNA fragment corresponding to *eipA* lacking the signal peptide (residues 39 - 198) was first PCR amplified. The primers used are listed in Table S6. The resulting PCR product was digested with BamHI / XhoI restriction enzymes (New England Biolabs) and ligated (T4 DNA ligase, New England Biolabs) into similarly digested pET28a plasmid. The ligation was then transformed into *E. coli* Top10 strain. After plating on LB agar plates + 50 μg ml^-1^ kanamycin, the resulting clones were sequence verified and transformed into the protein overexpression *E. coli* Rosetta(DE3) *plysS* strain (see Table S7 for strain information). For overexpression of His-tagged EipA, a 100-ml overnight Luria broth (LB; Fisher) culture (+ 50 μg ml^-1^ kanamycin) was inoculated with the protein expression strain and used to start a 1-liter LB flask (+ 50 μg ml^-1^ kanamycin) culture. Protein overexpression was induced at an OD600 ≈0.8 (37°C, 220 rpm) by adding 1 mM IPTG. After a 5-hour induction, cells were harvested by centrifugation at 11,000 × *g* for 20 min at 4°C. Cell pellets were resuspended in 25 ml of BugBuster Master Mix (Millipore - Novagen) and incubated for 20 minutes on ice. The resulting cell lysate was clarified by centrifugation at 39,000 x *g* for 20 min at 4°C. Purification of the His-tagged protein was performed using nickel affinity chromatography (nitrilotriacetic acid [NTA] resin; GE Healthcare). After binding of the clarified lysate samples to the column, two washing steps were performed, using Tris-NaCl buffers (10 mM Tris (pH 7.4) and 150 mM NaCl) containing 10 mM and 75 mM imidazole, followed by elution with Tris-NaCl buffer containing 500 mM imidazole. The protein purity of the different fractions was assessed by 14% SDS-PAGE and staining with Coomassie blue. Pure fractions were pooled and dialyzed against Tris-NaCl buffer (10 mM Tris (pH 8), 150 mM NaCl, 1 mM EDTA). EipA oligomeric state was then analyzed by size exclusion chromatography. Using a centrifugal filter unit (3-kDa molecular mass cutoff; Amicon-Millipore), a concentrated protein sample (500 μl at 5 mg ml^-1^) was injected onto a GE Healthcare Superdex 200 10/300 GL column (flow rate: 0.5 ml/min). An elution profile was measured at 280 nm and fractions of 500 μl were collected during the run; the dialysis buffer described above was used for all runs. Protein standards (blue dextran: 2,000 kDa, aldolase: 157 kDa, conalbumin: 76 kDa, and ovalbumin: 43 kDa) were also injected on to the column, and the corresponding calibration curve was use for molecular size estimation of the purified EipA.

### Bacterial two-hybrid protein interaction assay

To assay EipA self-interaction, we used a bacterial two-hybrid system (Battesti & Bouveret, 2012). *eipA* DNA fragments were cloned into pUT18 and pUT18C vector, which generated a C-terminal or N-terminal fusion to the T18 fragment of adenylate cyclase. *EipA* was also cloned into the pKT25 vector, which generated an N-terminal fusion to the T25 fragment of adenylate cyclase.

Primers used for PCRs are listed in Table S6 in the supplemental material; all plasmids were sequence confirmed. The different pUT18, pUT18C and pKT25 combinations were then cotransformed into chemically competent *E. coli* reporter strain BTH101 and plated on LB agar supplemented with ampicillin (100 μg ml^-1^) and kanamycin (50 μg ml^-1^). Control strains transformed with pKT25-Zip + pUT18C-Zip (positive control), and pKT25 + pUT18C (negative control), were also used. For each combination, a 3 ml LB culture tube containing the appropriate antibiotics was inoculated with a single colony and grown overnight at 30°C. The next day, 5 μl of each culture was spotted on an LB agar plate supplemented with ampicillin (100 μg ml^-1^), kanamycin (50 μg ml^-1^), and 5-bromo-4-chloro-3-indolyl-β-D-galactopyranoside (X-Gal, 40 μg ml^- 1^) and incubated at 30°C overnight. The next morning, positive interactions were confirmed by the blue color change of the spotted cultures.

### *Brucella* EipA overexpression strains

To ectopically express the different versions of EipA (wild-type, and EipA-PhoAEc fusion with or without the signal peptide) in *B. ovis*, the pSRKKm (Kan^R^) IPTG inducible plasmid was used (Khan *et al.*, 2008). An overlapping PCR strategy was used to stitch the *eipA* fragments (with or without the sequence corresponding to the signal peptide) to the *E. coli* alkaline phosphatase gene *phoA* (lacking its signal peptide) (primers are listed in Table S6). A Gibson-assembly (New England Biolabs) cloning strategy was then used to insert the different DNA fragments in the linearized pSRK plasmid. After sequencing, the different plasmids were introduced in *B. ovis* by overnight mating with *E. coli* WM3064 in presence of 300 μM of DAP and plated on SBA plates supplemented with kanamycin. See Tables S6 and S7 for primer and strain information.

### Alkaline phosphatase assay

To determine the cellular localization of EipA, we used a *B. ovis* strain transformed with the pSRK plasmid carrying *eipA* C-terminally fused to *E. coli phoA* (see Table S7). Two versions of this plasmid were made: one carrying the full-length *eipA*, which expressed the protein with its signal peptide, and one carrying a short version of *eipA*, which expressed the protein lacking the signal peptide. These two strains were grown overnight in 5 ml of Brucella broth supplemented with kanamycin (50 μg ml^-1^) and with or without 1 mM IPTG, allowing overexpression of the different protein fusions. The next day, the ODs (at 600 nm) were adjusted to ≈1.8 and 100 μl of each sample was transferred in a 96-well plate containing BCIP (final concentration: 200 μg ml^-1^). The same experiment was performed with spent medium supernatants. For each culture condition: 500 μl were centrifuged for 2 min at 14,000 rpm and 100 μl of the supernatants were transferred in a 96-well plate with BCIP. After 2 hours of incubation at 37°C / 5% CO2, the color change was visually assessed and pictures were taken. This experiment was conducted twice with two different clones each time.

### *B. ovis eipA* depletion assays

A *B. ovis* Δ*eipA* strain carrying plasmid pSRK-*eipA* (in which *eipA* expression is IPTG inducible) was used to evaluate the effect of depleting expression of *eipA*. *B. ovis* strains in which the chromosomal copy of *eipA* was intact, and that carried the empty pSRK vector or pSRK-*eipA*, were used as controls. 5 ml Brucella broth culture tubes were inoculated and grown overnight at 37°C with or without 1 mM IPTG. The next day, the optical densities (OD600) of the different cultures were adjusted to 0.15 (≈1 × 10^9^ CFU ml^-1^) in Brucella broth and serially diluted in a 96-well plate. 5 μl of each dilution was then spotted on a SBA plate (+ kanamycin 50 μg ml^-1^) supplemented (or not supplemented) with 1 mM IPTG. After three days of growth at 37°C / 5% CO2, colony forming units were compared between the different strains. Growth experiments in liquid were also performed with these strains. Overnight cultures were used to inoculate 5 ml of Brucella broth (+ kanamycin 50 μg ml^-1^) at a starting OD600 of 0.05. These cultures were incubated at 37°C / 5% CO2. For the first 12 hours, ODs were measured every 3 hours; a final OD was also taken after 24 hours of growth. Curve fit and doubling time calculations were performed using Prism 6 GraphPad software. Two different clones were used per strain and liquid growth measurements were performed in duplicate for each clone.

Phase-contrast images of cells from plates and liquid broth were collected using a Leica DM 5000B microscope with an HCX PL APO 63×/1.4 NA Ph3 objective. Images were acquired with a mounted Orca-ER digital camera (Hamamatsu) controlled by the Image-Pro software suite (Media Cybernetics). To prepare the different samples, colonies from SBA plates plus or minus 1 mM IPTG were resuspended in 1 ml PBS supplemented with 4% formaldehyde.

## Acknowledgments

We thank the members of the Crosson laboratory for helpful discussions. The authors wish to thank members of the SBC at Argonne National Laboratory for their help with data collection at the 19-ID beamline. This work was supported by National Institutes of Health Grants U19AI107792 and R01AI107159 to S.C.

## Author contributions

JH, JWW and SC contributed to the design and conceptualization of the study; JH, JWW, AF, LMV, DMC, JXC, EU, AB, LB, GB, YK and SC performed the experiments, acquired and analyzed the data; JH, JWW, AF and SC interpreted the data; JH and SC wrote the original draft of the manuscript.

## References

Adams, P.D., Grosse-Kunstleve, R.W., Hung, L.W., Ioerger, T.R., McCoy, A.J., Moriarty, N.W., et al. (2002) PHENIX: building new software for automated crystallographic structure determination. Acta Crystallogr D Biol Crystallogr 58: 1948–1954.

Alakomi, H.L., Skytta, E., Saarela, M., Mattila-Sandholm, T., Latva-Kala, K., and Helander, I.M. (2000) Lactic acid permeabilizes gram-negative bacteria by disrupting the outer membrane. Appl Environ Microbiol 66: 2001–2005.

Aslanidis, C., and de Jong, P.J. (1990) Ligation-independent cloning of PCR products (LIC-PCR). Nucleic Acids Res 18: 6069–6074.

Atluri, V.L., Xavier, M.N., de Jong, M.F., den Hartigh, A.B., and Tsolis, R.M. (2011) Interactions of the human pathogenic *Brucella* species with their hosts. Annu Rev Microbiol 65: 523–541.

Baldwin, C.L., and Parent, M. (2002) Fundamentals of host immune response against Brucella abortus: what the mouse model has revealed about control of infection. Vet Microbiol 90: 367–382.

Barnett, M.J., Hung, D.Y., Reisenauer, A., Shapiro, L., and Long, S.R. (2001) A homolog of the CtrA cell cycle regulator is present and essential in Sinorhizobium meliloti. J Bacteriol 183: 3204–3210.

Bateman, A., Coggill, P., and Finn, R.D. (2010) DUFs: families in search of function. Acta Crystallogr Sect F Struct Biol Cryst Commun 66: 1148–1152.

Battesti, A., and Bouveret, E. (2012) The bacterial two-hybrid system based on adenylate cyclase reconstitution in Escherichia coli. Methods 58: 325–334.

Batut, J., Andersson, S.G., and O’Callaghan, D. (2004) The evolution of chronic infection strategies in the alpha-proteobacteria. Nat Rev Microbiol 2: 933–945.

Bhavsar, A.P., Beveridge, T.J., and Brown, E.D. (2001) Precise deletion of tagD and controlled depletion of its product, glycerol 3-phosphate cytidylyltransferase, leads to irregular morphology and lysis of Bacillus subtilis grown at physiological temperature. J. Bacteriol. 183: 6688–6693.

Bochner, B.R. (2009) Global phenotypic characterization of bacteria. FEMS Microbiol Rev 33: 191–205.

Brilli, M., Fondi, M., Fani, R., Mengoni, A., Ferri, L., Bazzicalupo, M., et al. (2010) The diversity and evolution of cell cycle regulation in alpha-proteobacteria: a comparative genomic analysis. BMC Syst Biol 4: 52.

Bush, K. (2018) Past and present perspectives on beta-lactamases. Antimicrob. Agents Chemother. 62.

Byndloss, M.X., and Tsolis, R.M. (2016) Brucella spp. virulence factors and immunity. Annu Rev Anim Biosci 4: 111–127.

Cabeen, M.T., Murolo, M.A., Briegel, A., Bui, N.K., Vollmer, W., Ausmees, N., et al. (2010) Mutations in the lipopolysaccharide biosynthesis pathway interfere with crescentin-mediated cell curvature in Caulobacter crescentus. J Bacteriol 192: 3368–3378.

Cardoso, P.G., Macedo, G.C., Azevedo, V., and Oliveira, S.C. (2006) Brucella spp noncanonical LPS: structure, biosynthesis, and interaction with host immune system. Microb Cell Fact 5: 13.

Celli, J. (2006) Surviving inside a macrophage: the many ways of Brucella. Res Microbiol 157: 9398.

Cloeckaert, A., Grayon, M., Verger, J.M., Letesson, J.J., and Godfroid, F. (2000) Conservation of seven genes involved in the biosynthesis of the lipopolysaccharide O-side chain in Brucella spp. Res Microbiol 151: 209–216.

Conde-Alvarez, R., Arce-Gorvel, V., Iriarte, M., Mancek-Keber, M., Barquero-Calvo, E., Palacios-Chaves, L., et al. (2012) The lipopolysaccharide core of Brucella abortus acts as a shield against innate immunity recognition. PLoS Pathog 8: e1002675.

Dabral, N., Jain-Gupta, N., Seleem, M.N., Sriranganathan, N., and Vemulapalli, R. (2015) Overexpression of Brucella putative glycosyltransferase WbkA in B. abortus RB51 leads to production of exopolysaccharide. Front Cell Infect Microbiol 5: 54.

de Barsy, M., Jamet, A., Filopon, D., Nicolas, C., Laloux, G., Rual, J.F., et al. (2011) Identification of a Brucella spp. secreted effector specifically interacting with human small GTPase Rab2. Cell Microbiol 13: 1044–1058.

De Bolle, X., Crosson, S., Matroule, J.Y., and Letesson, J.J. (2015) Brucella abortus cell cycle and infection are coordinated. Trends Microbiol 23: 812–821.

de Figueiredo, P., Ficht, T.A., Rice-Ficht, A., Rossetti, C.A., and Adams, L.G. (2015) Pathogenesis and immunobiology of brucellosis: review of *Brucella*-host interactions. Am J Pathol 185: 1505–1517.

De Nisco, N.J., Abo, R.P., Wu, C.M., Penterman, J., and Walker, G.C. (2014) Global analysis of cell cycle gene expression of the legume symbiont Sinorhizobium meliloti. Proc Natl Acad Sci U S A 111: 3217–3224.

Delrue, R.M., Martinez-Lorenzo, M., Lestrate, P., Danese, I., Bielarz, V., Mertens, P., et al. (2001) Identification of Brucella spp. genes involved in intracellular trafficking. Cell Microbiol 3: 487–497.

den Hartigh, A.B., Rolan, H.G., de Jong, M.F., and Tsolis, R.M. (2008) VirB3 to VirB6 and VirB8 to VirB11, but not VirB7, are essential for mediating persistence of Brucella in the reticuloendothelial system. J Bacteriol 190: 4427–4436.

Duxbury, C.L., and Thompson, J.E. (1987) Pentachlorophenol alters the molecular organization of membranes in mammalian cells. Arch Environ Contam Toxicol 16: 367–373.

Emsley, P., and Cowtan, K. (2004) Coot: model-building tools for molecular graphics. Acta Crystallogr D Biol Crystallogr 60: 2126–2132.

Eschenfeldt, W.H., Lucy, S., Millard, C.S., Joachimiak, A., and Mark, I.D. (2009) A family of LIC vectors for high-throughput cloning and purification of proteins. Methods Mol Biol 498: 105–115.

Fang, G., Passalacqua, K.D., Hocking, J., Llopis, P.M., Gerstein, M., Bergman, N.H., et al. (2013) Transcriptomic and phylogenetic analysis of a bacterial cell cycle reveals strong associations between gene co-expression and evolution. BMC Genomics 14: 450.

Finn, R.D., Coggill, P., Eberhardt, R.Y., Eddy, S.R., Mistry, J., Mitchell, A.L., et al. (2016) The Pfam protein families database: towards a more sustainable future. Nucleic Acids Res 44: D279–285.

Francis, N., Poncin, K., Fioravanti, A., Vassen, V., Willemart, K., Ong, T.A., et al. (2017) CtrA controls cell division and outer membrane composition of the pathogen Brucella abortus. Mol Microbiol 103: 780–797.

Gonzalez, D., Grillo, M.J., De Miguel, M.J., Ali, T., Arce-Gorvel, V., Delrue, R.M., et al. (2008) Brucellosis vaccines: assessment of Brucella melitensis lipopolysaccharide rough mutants defective in core and O-polysaccharide synthesis and export. PLoS One 3: e2760.

Gorvel, J.P. (2008) Brucella: a Mr “Hide” converted into Dr Jekyll. Microbes Infect 10: 1010–1013.

Gorvel, J.P., and Moreno, E. (2002) Brucella intracellular life: from invasion to intracellular replication. Vet Microbiol 90: 281–297.

Guzman-Verri, C., Manterola, L., Sola-Landa, A., Parra, A., Cloeckaert, A., Garin, J., et al. (2002) The two-component system BvrR/BvrS essential for Brucella abortus virulence regulates the expression of outer membrane proteins with counterparts in members of the Rhizobiaceae. Proc Natl Acad Sci U S A 99: 12375–12380.

Hagemans, D., van Belzen, I.A., Moran Luengo, T., and Rudiger, S.G. (2015) A script to highlight hydrophobicity and charge on protein surfaces. Front Mol Biosci 2: 56.

Heneberg, P. (2012) Finding the smoking gun: protein tyrosine phosphatases as tools and targets of unicellular microorganisms and viruses. Curr Med Chem 19: 1530–1566.

Holm, L., and Laakso, L.M. (2016) Dali server update. Nucleic Acids Res 44: W351–355.

Holm, L., and Rosenstrom, P. (2010) Dali server: conservation mapping in 3D. Nucleic Acids Res 38: W545–549.

Hottes, A.K., Shapiro, L., and McAdams, H.H. (2005) DnaA coordinates replication initiation and cell cycle transcription in Caulobacter crescentus. Mol Microbiol 58: 1340–1353.

Jorgenson, M.A., and Young, K.D. (2016) Interrupting biosynthesis of O-antigen or the lipopolysaccharide core produces morphological defects in Escherichia coli by sequestering undecaprenyl phosphate. J. Bacteriol. 198: 3070–3079.

Kainth, P., and Gupta, R.S. (2005) Signature proteins that are distinctive of alpha proteobacteria. BMC Genomics 6: 94.

Khan, S.R., Gaines, J., Roop, R.M., 2nd, and Farrand, S.K. (2008) Broad-host-range expression vectors with tightly regulated promoters and their use to examine the influence of TraR and TraM expression on Ti plasmid quorum sensing. Appl Environ Microbiol 74: 5053–5062.

Kim, H.S., Caswell, C.C., Foreman, R., Roop, R.M., 2nd, and Crosson, S. (2013) The Brucella abortus general stress response system regulates chronic mammalian infection and is controlled by phosphorylation and proteolysis. J Biol Chem 288: 13906–13916.

Kim, Y., Babnigg, G., Jedrzejczak, R., Eschenfeldt, W.H., Li, H., Maltseva, N., et al. (2011) High-throughput protein purification and quality assessment for crystallization. Methods 55: 12–28.

Lamontagne, J., Butler, H., Chaves-Olarte, E., Hunter, J., Schirm, M., Paquet, C., et al. (2007) Extensive cell envelope modulation is associated with virulence in Brucella abortus. J Proteome Res 6: 1519–1529.

Lapaque, N., Moriyon, I., Moreno, E., and Gorvel, J.P. (2005) Brucella lipopolysaccharide acts as a virulence factor. Curr Opin Microbiol 8: 60–66.

Lasker, K., Schrader, J.M., Men, Y., Marshik, T., Dill, D.L., McAdams, H.H., et al. (2016) CauloBrowser: A systems biology resource for Caulobacter crescentus. Nucleic Acids Res. 44: D640–645.

Laskowski, R.A., MacArthur, M.W., Moss, D.S., and Thornton, J.M. (1993) PROCHECK: a program to check the stereochemical quality of protein structures. J Appl Crystallogr 26: 283–291.

Laub, M.T., Chen, S.L., Shapiro, L., and McAdams, H.H. (2002) Genes directly controlled by CtrA, a master regulator of the Caulobacter cell cycle. Proc Natl Acad Sci U S A 99: 4632–4637.

Laub, M.T., McAdams, H.H., Feldblyum, T., Fraser, C.M., and Shapiro, L. (2000) Global analysis of the genetic network controlling a bacterial cell cycle. Science 290: 2144–2148.

Manat, G., Roure, S., Auger, R., Bouhss, A., Barreteau, H., Mengin-Lecreulx, D., et al. (2014) Deciphering the metabolism of undecaprenyl-phosphate: the bacterial cell-wall unit carrier at the membrane frontier. Microb. Drug Resist. 20: 199–214.

Mancilla, M. (2015) Smooth to rough dissociation in *Brucella*: the missing link to virulence. Front Cell Infect Microbiol 5: 98.

Mancilla, M., Marin, C.M., Blasco, J.M., Zarraga, A.M., Lopez-Goni, I., and Moriyon, I. (2012) Spontaneous excision of the O-polysaccharide wbkA glycosyltranferase gene is a cause of dissociation of smooth to rough Brucella colonies. J Bacteriol 194: 1860–1867.

Manterola, L., Moriyon, I., Moreno, E., Sola-Landa, A., Weiss, D.S., Koch, M.H., et al. (2005) The lipopolysaccharide of Brucella abortus BvrS/BvrR mutants contains lipid A modifications and has higher affinity for bactericidal cationic peptides. J Bacteriol 187: 5631–5639.

Marrichi, M., Camacho, L., Russell, D.G., and DeLisa, M.P. (2008) Genetic toggling of alkaline phosphatase folding reveals signal peptides for all major modes of transport across the inner membrane of bacteria. J Biol Chem 283: 35223–35235.

Martinez de Tejada, G., Pizarro-Cerda, J., Moreno, E., and Moriyon, I. (1995) The outer membranes of Brucella spp. are resistant to bactericidal cationic peptides. Infect. Immun. 63: 3054–3061.

McDonnell, G., and Russell, A.D. (1999) Antiseptics and disinfectants: activity, action, and resistance. Clin Microbiol Rev 12: 147–179.

McGrath, P.T., Iniesta, A.A., Ryan, K.R., Shapiro, L., and McAdams, H.H. (2006) A dynamically localized protease complex and a polar specificity factor control a cell cycle master regulator. Cell 124: 535–547.

McGrath, P.T., Lee, H., Zhang, L., Iniesta, A.A., Hottes, A.K., Tan, M.H., et al. (2007) High-throughput identification of transcription start sites, conserved promoter motifs and predicted regulons. Nat Biotechnol 25: 584–592.

Minor, W., Cymborowski, M., Otwinowski, Z., and Chruszcz, M. (2006) HKL-3000: the integration of data reduction and structure solution--from diffraction images to an initial model in minutes. Acta Crystallogr D Biol Crystallogr 62: 859–866.

Monreal, D., Grillo, M.J., Gonzalez, D., Marin, C.M., De Miguel, M.J., Lopez-Goni, I., et al. (2003) Characterization of Brucella abortus O-polysaccharide and core lipopolysaccharide mutants and demonstration that a complete core is required for rough vaccines to be efficient against Brucella abortus and Brucella ovis in the mouse model. Infect Immun 71: 3261–3271.

Murshudov, G.N., Vagin, A.A., and Dodson, E.J. (1997) Refinement of macromolecular structures by the maximum-likelihood method. Acta Crystallogr D Biol Crystallogr 53: 240–255.

Myeni, S., Child, R., Ng, T.W., Kupko, J.J., 3rd, Wehrly, T.D., Porcella, S.F., et al. (2013) Brucella modulates secretory trafficking via multiple type IV secretion effector proteins. PLoS Path. 9: e1003556.

Nielsen, H. (2017) Predicting Secretory Proteins with SignalP. Methods Mol Biol 1611: 59–73.

O’Callaghan, D., Cazevieille, C., Allardet-Servent, A., Boschiroli, M.L., Bourg, G., Foulongne, V., et al. (1999) A homologue of the Agrobacterium tumefaciens VirB and Bordetella pertussis Ptl type IV secretion systems is essential for intracellular survival of Brucella suis. Mol Microbiol 33: 1210–1220.

Oetken, C., von Willebrand, M., Autero, M., Ruutu, T., Andersson, L.C., and Mustelin, T. (1992) Phenylarsine oxide augments tyrosine phosphorylation in hematopoietic cells. Eur J Haematol 49: 208–214.

Ohsuka, S., Ohta, M., Masuda, K., Arakawa, Y., Kaneda, T., and Kato, N. (1994) Lidocaine hydrochloride and acetylsalicylate kill bacteria by disrupting the bacterial membrane potential in different ways. Microbiol Immunol 38: 429–434.

Okoko, T., Blagova, E.V., Whittingham, J.L., Dover, L.G., and Wilkinson, A.J. (2015) Structural characterisation of the virulence-associated protein VapG from the horse pathogen Rhodococcus equi. Vet Microbiol 179: 42–52.

Oliveira, S.C., de Almeida, L.A., Carvalho, N.B., Oliveira, F.S., and Lacerda, T.L. (2012) Update on the role of innate immune receptors during Brucella abortus infection. Vet Immunol Immunopathol 148: 129–135.

Palmer, D.A., and Douglas, J.T. (1989) Analysis of Brucella lipopolysaccharide with specific and cross-reacting monoclonal antibodies. J. Clin. Microbiol. 27: 2331–2337.

Panis, G., Murray, S.R., and Viollier, P.H. (2015) Versatility of global transcriptional regulators in alpha-proteobacteria: from essential cell cycle control to ancillary functions. FEMS Microbiol Rev 39: 120–133.

Pei, J., Ding, X., Fan, Y., Rice-Ficht, A., and Ficht, T.A. (2012) Toll-like receptors are critical for clearance of Brucella and play different roles in development of adaptive immunity following aerosol challenge in mice. Front Cell Infect Microbiol 2: 115.

Perry, B.J., Akter, M.S., and Yost, C.K. (2016) The use of transposon insertion sequencing to interrogate the core functional genome of the legume symbiont Rhizobium leguminosarum. Front Microbiol 7: 1873.

Pini, F., De Nisco, N.J., Ferri, L., Penterman, J., Fioravanti, A., Brilli, M., et al. (2015) Cell cycle control by the master regulator CtrA in Sinorhizobium meliloti. PLoS Genet 11: e1005232.

Poncin, K., Gillet, S., and De Bolle, X. (2018) Learning from the master: targets and functions of the CtrA response regulator in Brucella abortus and other alpha-proteobacteria. FEMS Microbiol Rev 42: 500–513.

Price, M.N., Wetmore, K.M., Waters, R.J., Callaghan, M., Ray, J., Liu, H., et al. (2018) Mutant phenotypes for thousands of bacterial genes of unknown function. Nature 557: 503–509.

Reisenauer, A., Quon, K., and Shapiro, L. (1999) The CtrA response regulator mediates temporal control of gene expression during the Caulobacter cell cycle. J. Bacteriol. 181: 2430–2439.

Rick, P.D., Barr, K., Sankaran, K., Kajimura, J., Rush, J.S., and Waechter, C.J. (2003) Evidence that the wzxE gene of Escherichia coli K-12 encodes a protein involved in the transbilayer movement of a trisaccharide-lipid intermediate in the assembly of enterobacterial common antigen. J. Biol. Chem. 278: 16534–16542.

Salvador-Bescos, M., Gil-Ramirez, Y., Zuniga-Ripa, A., Martinez-Gomez, E., de Miguel, M.J., Munoz, P.M., et al. (2018) WadD, a new Brucella lipopolysaccharide core glycosyltransferase identified by genomic search and phenotypic characterization. Front. Microbiol. 9: 2293.

Santa Maria, J.P., Jr., Sadaka, A., Moussa, S.H., Brown, S., Zhang, Y.J., Rubin, E.J., et al. (2014) Compound-gene interaction mapping reveals distinct roles for Staphylococcus aureus teichoic acids. Proc. Natl. Acad. Sci. U. S. A. 111: 12510–12515.

Scariot, F.J., Jahn, L., Delamare, A.P.L., and Echeverrigaray, S. (2017) Necrotic and apoptotic cell death induced by Captan on Saccharomyces cerevisiae. World J Microbiol Biotechnol 33: 159.

Schluter, J.P., Reinkensmeier, J., Barnett, M.J., Lang, C., Krol, E., Giegerich, R., et al. (2013) Global mapping of transcription start sites and promoter motifs in the symbiotic alpha-proteobacterium Sinorhizobium meliloti 1021. BMC Genomics 14: 156.

Schrader, J.M., Li, G.W., Childers, W.S., Perez, A.M., Weissman, J.S., Shapiro, L., et al. (2016) Dynamic translation regulation in Caulobacter cell cycle control. Proc Natl Acad Sci U S A 113: E6859–E6867.

Siam, R., and Marczynski, G.T. (2000) Cell cycle regulator phosphorylation stimulates two distinct modes of binding at a chromosome replication origin. EMBO J. 19: 1138–1147.

Slabinski, L., Jaroszewski, L., Rychlewski, L., Wilson, I.A., Lesley, S.A., and Godzik, A. (2007) XtalPred: a web server for prediction of protein crystallizability. Bioinformatics 23: 3403–3405.

Smith, J.A. (2018) Brucella lipopolysaccharide and pathogenicity: the core of the matter. Virulence 9: 379–382.

Smith, S.C., Joshi, K.K., Zik, J.J., Trinh, K., Kamajaya, A., Chien, P., et al. (2014) Cell cycle-dependent adaptor complex for ClpXP-mediated proteolysis directly integrates phosphorylation and second messenger signals. Proc Natl Acad Sci U S A 111: 14229–14234.

Soler-Llorens, P., Gil-Ramirez, Y., Zabalza-Barangua, A., Iriarte, M., Conde-Alvarez, R., Zuniga-Ripa, A., et al. (2014) Mutants in the lipopolysaccharide of Brucella ovis are attenuated and protect against B. ovis infection in mice. Vet Res 45: 72.

Spencer, W., Siam, R., Ouimet, M.C., Bastedo, D.P., and Marczynski, G.T. (2009) CtrA, a global response regulator, uses a distinct second category of weak DNA binding sites for cell cycle transcription control in Caulobacter crescentus. J Bacteriol 191: 5458–5470.

Spera, J.M., Herrmann, C.K., Roset, M.S., Comerci, D.J., and Ugalde, J.E. (2013) A Brucella virulence factor targets macrophages to trigger B-cell proliferation. J Biol Chem 288: 20208–20216.

Standish, A.J., and Morona, R. (2014) The role of bacterial protein tyrosine phosphatases in the regulation of the biosynthesis of secreted polysaccharides. Antioxid Redox Signal 20: 2274–2289.

Tatar, L.D., Marolda, C.L., Polischuk, A.N., van Leeuwen, D., and Valvano, M.A. (2007) An Escherichia coli undecaprenyl-pyrophosphate phosphatase implicated in undecaprenyl phosphate recycling. Microbiology 153: 2518–2529.

Tian, M., Qu, J., Han, X., Ding, C., Wang, S., Peng, D., et al. (2014) Mechanism of Asp24 upregulation in Brucella abortus rough mutant with a disrupted O-antigen export system and effect of Asp24 in bacterial intracellular survival. Infect Immun 82: 2840–2850.

Tsolis, R.M., Seshadri, R., Santos, R.L., Sangari, F.J., Lobo, J.M., de Jong, M.F., et al. (2009) Genome degradation in Brucella ovis corresponds with narrowing of its host range and tissue tropism. PLoS One 4: e5519.

Ugalde, J.E., Czibener, C., Feldman, M.F., and Ugalde, R.A. (2000) Identification and characterization of the Brucella abortus phosphoglucomutase gene: role of lipopolysaccharide in virulence and intracellular multiplication. Infect Immun 68: 5716–5723.

Wallace, I.M., O’Sullivan, O., Higgins, D.G., and Notredame, C. (2006) M-Coffee: combining multiple sequence alignment methods with T-Coffee. Nucleic Acids Res 34: 1692–1699.

Weiss, D.S., Takeda, K., Akira, S., Zychlinsky, A., and Moreno, E. (2005) MyD88, but not toll-like receptors 4 and 2, is required for efficient clearance of Brucella abortus. Infect Immun 73: 5137–5143.

Wetmore, K.M., Price, M.N., Waters, R.J., Lamson, J.S., He, J., Hoover, C.A., et al. (2015) Rapid quantification of mutant fitness in diverse bacteria by sequencing randomly bar-coded transposons. MBio 6: e00306–00315.

Weynants, V., Gilson, D., Cloeckaert, A., Denoel, P.A., Tibor, A., Thiange, P., et al. (1996) Characterization of a monoclonal antibody specific for Brucella smooth lipopolysaccharide and development of a competitive enzyme-linked immunosorbent assay to improve the serological diagnosis of brucellosis. Clin. Vaccine Immunol. 3: 309–314.

Willett, J.W., Herrou, J., Briegel, A., Rotskoff, G., and Crosson, S. (2015) Structural asymmetry in a conserved signaling system that regulates division, replication, and virulence of an intracellular pathogen. Proc Natl Acad Sci U S A 112: E3709–3718.

Williams, K.P., Sobral, B.W., and Dickerman, A.W. (2007) A robust species tree for the alphaproteobacteria. J. Bacteriol. 189: 4578–4586.

Yuasa, R., Levinthal, M., and Nikaido, H. (1969) Biosynthesis of cell wall lipopolysaccharide in mutants of Salmonella. V. A mutant of Salmonella typhimurium defective in the synthesis of cytidine diphosphoabequose. J. Bacteriol. 100: 433–444.

Zhang, M., Han, X., Liu, H., Tian, M., Ding, C., Song, J., et al. (2013) Inactivation of the ABC transporter ATPase gene in Brucella abortus strain 2308 attenuated the virulence of the bacteria. Vet Microbiol 164: 322–329.

Zhou, B., Schrader, J.M., Kalogeraki, V.S., Abeliuk, E., Dinh, C.B., Pham, J.Q., et al. (2015) The global regulatory architecture of transcription during the Caulobacter cell cycle. PLoS Genet 11: e1004831.

Zygmunt, M.S., Blasco, J.M., Letesson, J.J., Cloeckaert, A., and Moriyon, I. (2009) DNA polymorphism analysis of *Brucella* lipopolysaccharide genes reveals marked differences in O-polysaccharide biosynthetic genes between smooth and rough Brucella species and novel species-specific markers. BMC Microbiol 9: 92.

